# A histone-like motif in yellow fever virus contributes to viral replication

**DOI:** 10.1101/2020.05.05.078782

**Authors:** Diego Mourão, Shoudeng Chen, Uwe Schaefer, Leonia Bozzacco, Leticia A. Carneiro, Alan Gerber, Carolina Adura, Brian D. Dill, Henrik Molina, Thomas Carroll, Matthew Paul, Natarajan V. Bhanu, Benjamin A. Garcia, Gerard Joberty, Inmaculada Rioja, Rab K. Prinjha, Robert G. Roeder, Charles M. Rice, Margaret R. MacDonald, Dinshaw Patel, Alexander Tarakhovsky

## Abstract

The mimicry of host proteins by viruses contributes to their ability to suppress antiviral immunity and hijack host biosynthetic machinery^1^. Host adaptation to evade this exploitation depends on host protein functional redundancy^2^. Non-redundant, essential host proteins have limited potential to adapt without severe consequences^3^. Histones, which are essential for genome architecture and control of gene expression, are among the most evolutionary conserved proteins^4^. Here we show that the capsid protein of the flavivirus yellow fever virus (YFV), mimics histone H4 and interferes with chromatin gene regulation by BRD4, a bromodomain and extraterminal domain (BET) protein. Two acetyl-lysine residues of YFV capsid are embedded in a histone-like motif that interacts with the BRD4 bromodomain, affecting gene expression and influencing YFV replication. These findings reveal histone mimicry as a strategy employed by an RNA virus that replicates in the cytosol^5^ and define convergent and distinct molecular determinants for motif recognition of the viral mimic versus histone H4.

---

Flaviviruses, like YFV, are positive-sense RNA viruses that replicate in the cytosol in association with modified cellular membranes^5^. Despite that, some flavivirus proteins such as NS5 and capsid protein (C) traffic to the nucleus where they presumably perform additional functions besides RNA replication (as for NS5) or viral genome encapsidation (as for C)^5,6^. A fraction of flavivirus C, as for dengue, accumulates in the nucleus and nucleolus of the infected cells, where C can bind to nucleosomes or individual histones^7,8^. Indeed, analysis of capsid sub-cellular distribution upon YFV infection of the human hepatoma cell line Huh-7 shows that YFV capsid (hereafter referred as YFV-C) is found at similar levels both in the nucleus and cytosol of infected cells (Fig. 1a). Moreover, nuclear YFV-C accumulates in the nucleolus of infected cells (Fig. 1b) as previously reported for other flaviviruses^7^. The potential impact of YFV-C on nuclear function is suggested by the high abundance of YFV-C in the nuclei of the infected cells. Using purified YFV-C as a standard we determined the amount of YFV-C per nucleus of cells at different time points after infection (Extended Data Fig. 1a). According to our calculations, we estimate that approximately 10^7^ molecules of YFV-C (around 50% of the total) are present in the nucleus of cells 24 hr after infection (Extended Data Fig. 1a,b). Moreover, YFV-C is tightly bound to chromatin as assessed by NUN (NaCl, Urea, NP-40) extraction procedure of YFV-C from the nuclei of YFV-infected Huh-7 (Fig. 1c), which leaves tightly bound proteins (mainly histones) in the chromatin pellet^9^.

**Figure 1.**
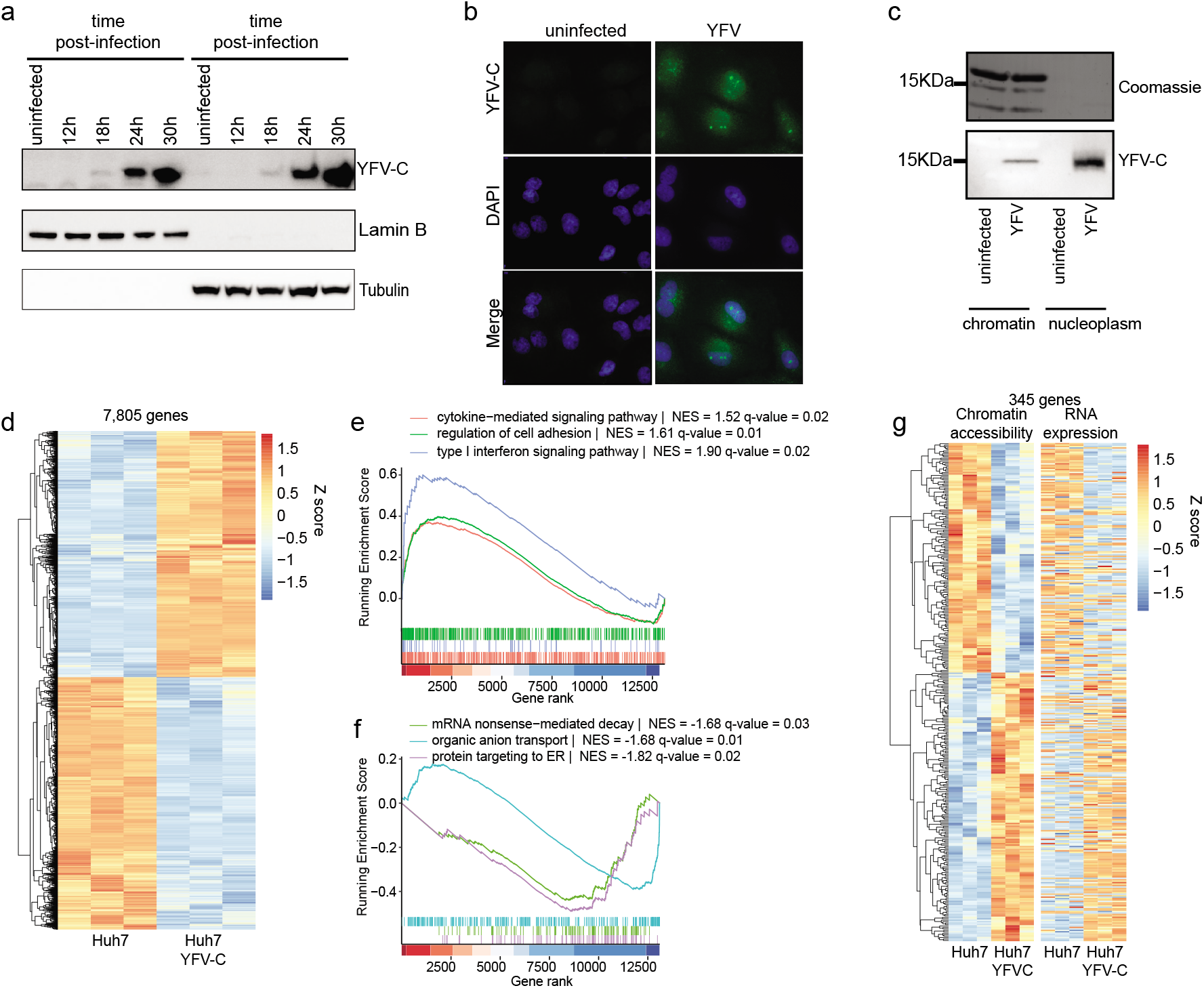
Capsid protein from yellow fever virus is a chromatin-binding protein that modulates gene expression and chromatin accessibility. **a,** Huh-7 cells were mock infected or infected with YFV 17D (multiplicity of infection (MOI)=10) for the indicated time upon which cells were harvested and fractionated into cytosolic and nuclear fractions. Protein lysates were analyzed by western blot using a newly developed rabbit anti-YFV-C polyclonal antibody as well as anti-α-Tubulin and anti-Lamin B. Data is representative of at least 3 independent experiments. **b**, Huh-7 cells were infected with YFV 17D (MOI=10) for 24 hours prior to fixation and immunofluorescence staining with rabbit anti-YFV-C and DAPI. Cells were analyzed using a confocal microscope. Data is representative of 3 independent experiments. **c,** Huh-7 cells were infected with YFV 17D (MOI=10) for 24 hr prior to cell nuclear and cytosolic detergent fractionation. The obtained nuclei were further fractionated into nucleoplasm and chromatin fractions using NUN buffer. Chromatin-bound proteins were eluted by incubation at 40°C in 8M urea-containing buffer followed by sonication. **d**, RNA isolated from Huh-7 cells expressing an empty vector (Huh-7) or expressing YFV-C for 48h (Huh-7 YFV-C) were sequenced and analyzed for gene expression changes. Genes with an adjusted p-value < 0.001 were considered differentially expressed (7,805 genes) and were plotted in a heatmap. Data is representative of 3 technical replicates. **e,f,** All expressed RNA were ordered by rank of gene expression differences between Huh-7 YFV-C vs Huh-7. The ranked list was analyzed by GSEA for suppressed (e) or induced (f) GO biological process terms. **g,** Nuclei isolated from Huh-7 cells expressing an empty vector (Huh-7) or expressing YFV-C for 48h (Huh-7 YFV-C) were processed and chromatin accessibility was quantified. Genes showing chromatin accessibility changes and corresponding expressed RNA (345 genes) were plotted in a heatmap. Data is representative of 3 technical replicates.

Nuclear compartmentalization and chromatin interaction of YFV-C can trigger major changes in chromatin and gene expression. We found that ectopically expressed YFV-C, reaching around 10% of the nuclear YFV-C levels of 24 hr YFV-infected Huh-7 cells (Extended Data Fig. 1c,d), is associated with genome-wide gene expression changes as assessed by RNA-seq assay. Expression of YFV-C for 48 hr (designated Huh-7 YFV-C) causes changes in 7,805 genes (adjusted p-value < 0.001) with down- or up-regulation of 3,915 or 3,890 genes, respectively, in comparison with Huh-7 cells with an empty vector (Huh-7) (Fig. 1d). Gene set enrichment analysis (GSEA) showed that organic anion transport (which includes cholesterol and sterol transport), protein targeting to endoplasmic reticulum (ER) and mRNA nonsense-mediated decay are among the biological processes with the most down-regulated genes, while up-regulated genes are involved on cell adhesion, cytokine-mediated signaling and type I interferon signaling (Fig. 1e and Extended Data Fig. 2a). Among the suppressed biological processes, rRNA metabolic process and nonsense-mediated decay were already reported to be targeted by other flaviviruses^10–12^, while induction of cytokine-mediated signaling and type I interferon signaling are known responses to flavivirus infection^13^.

The presence of a tightly bound fraction of YFV-C in the chromatin suggests a capacity to modulate chromatin accessibility. Quantification of genome-wide changes in chromatin as measured by Assay for Accessible Chromatin sequencing (ATAC-seq) showed 2,970 accessible loci, with the majority of accessible loci located at the transcription start site (TSS) of genes (Extended Data Fig. 2b). Expression of YFV-C increased and decreased chromatin accessibility in 271 and 203 loci respectively, without any bias in the genomic localization (Extended Data Fig. 2c,d). The effect of YFV-C on chromatin partially correlates with gene expression changes caused by YFV-C (Fig. 1g and Extended Data Fig. 2e-g).

We wondered how YFV-C might exert these effects on gene expression. We noted that YFV-C displays between 25-37% similarity to canonical histone proteins (Extended Data Fig. 3a-d) and bears 75% similarity with N-terminal unstructured segments of the histone H4 (Fig. 2a). In particular, YFV-C contains a short linear motif (KAQGK) that is similar to one of the acetylation motifs of the histone H4 (KGGK). This H4-like motif of YFV-C is well conserved among the 37 different YFV strains obtained from the Virus Pathogen Resource database (ViPR) ^14^ (Extended Data Fig. 3e) and lies in a long unstructured domain^15^.

**Figure 2.**
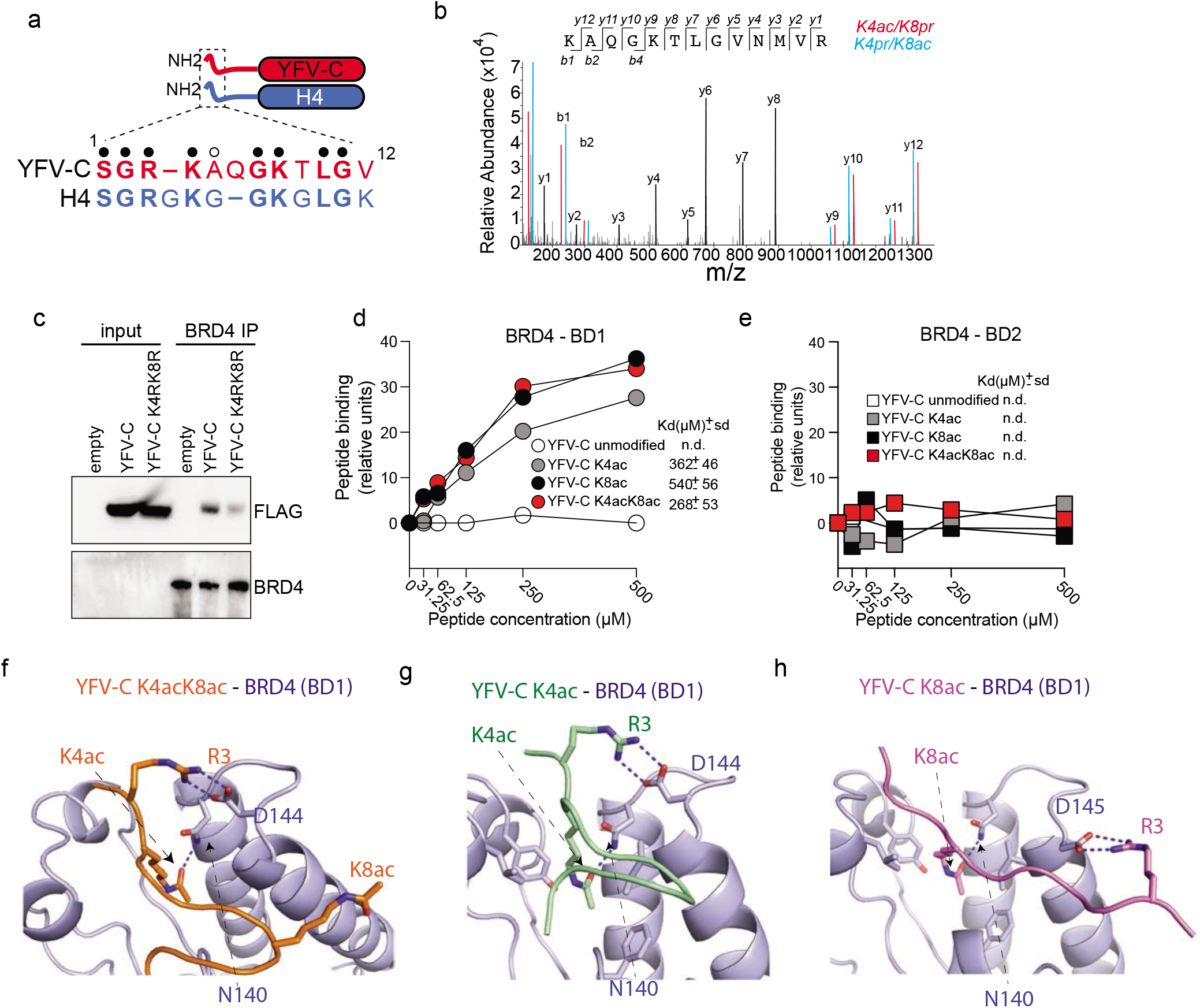
YFV-C protein possesses a histone H4-like acetylatable motif that binds BRD4. **a**, Graphical depiction of the unstructured N-terminal portion of YFV-C and H4 showing alignment of the 12 most N-terminal amino acids. Amino acid identity is depicted with a closed circle, amino acid side chain similarity is depicted with an open circle and gaps in the alignment are depicted by a dash. **b,** Huh-7 cells were infected for 18 hr with YFV 17D (MOI=1) prior to nuclei isolation and immunoprecipitation of YFV-C. The isolated material was separated in a polyacrylamide gel and the excised band was analyzed by LC-MS/MS to detect post-translational modifications of YFV-C. Lysine acetylation was detected in lysine 4 or lysine 8. The MS/MS chimeric spectrum of the tryptic peptide KAQGKTLGVNMVR from YFV-C protein (K4ac-K8pr in red and K4pr-K8ac in blue) is shown. Propionylation (pr) stems from the chemical derivatization of the digested protein. All ions detected were the prominent peaks well above the noise and are annotated with the mass/charge (m/z). **c,** T-Rex Flp-In HeLa cells carrying an empty vector (empty), vector expressing 3XFlag epitope-tagged YFV-C (WT) or 3XFlag tagged YFV-C with K4 to R and K8 to R mutations (K4RK8R) were treated with 1μg/ml of doxycycline for 24 hours for YFV-C gene expression induction prior to cell lysis and BRD4 immunoprecipitation, polyacrylamide gel separation and western blot probing for BRD4 and Flag as indicated. 1% of the input material is shown (left) as well as the immunoprecipitated material (right). Data is representative of 3 independent experiments. **d,e,** Surface plasmon resonance (SPR) equilibrium plots of interactions of lysine acetylated YFV-C (residues 1-11) with bromodomain 1 (d) and bromodomain 2 (e) of BRD4. The constant of dissociation (Kd) was calculated based on the equilibrium between 45 and 50 seconds. Data is representative of at least 3 independent experiments. **f-h,** Intermolecular interactions within and in the vicinity of the bromodomain binding pocket. The residues Arginine 3 (R3), Lysine 4 acetylated (K4ac), Lysine 8 acetylated (K8ac) of YFV-C protein and Asparagine 140 (N140), Aspartic acid 144 (D144) and 145 (D145) of Brd4 are highlighted with stick representations in K4acK8ac (f), K4ac (g) and K8ac (h). Brd4 is shown in light blue, while YFV-C residues are colored as indicated.

We wondered whether similar to the histone H4, lysine 4 or 8 of YFV-C can be acetylated in the nucleus upon virus infection. The MS/MS analysis of YFV-C isolated from Huh-7 cells infected with the 17D strain of YFV for 18 hours shows the presence of acetylated lysine 4 or 8 (Fig. 2b) and di-acetylated lysine 4 and 8 (Extended Data Fig. 4a). Acetylation of YFV-C does not require infection and could be detected in YFV-C ectopically expressed for 24 hr in HeLa cells (Extended Data Fig. 4b,c). The presence of a histone H4-like motif in YFV-C and its acetylation underscores the histone H4 mimicry by YFV-C.

Acetylation of lysine residues on histone H4 N-terminus plays a key role in gene expression^16^. Much of this histone H4 function is mediated by the bromodomain containing BET proteins that bind selectively to acetylated lysine residues at the histone H4 N-terminus^17,18^. We found that BRD4 immunoprecipitation from HeLa cells leads to co-precipitation of YFV-C. Substitution of lysine 4 and lysine 8 to arginine (K4RK8R) reduced the abundance of YFV-C associated with BRD4 (Fig. 2c and Extended Data Fig. 5a), BRD3 and BRD2 (Extended Data Fig. 5b-e). Diminished interaction of YFV-C K4RK8R with BET proteins is not due to differences in subcellular distribution as it is found at similar levels to YFV-C in the nucleus and chromatin (Extended Data Fig. 6).

Direct interaction assay by surface plasmon resonance confirmed that YFV-C histone mimic-containing peptide associates with bromodomain 1, but not bromodomain 2, of BRD4 in an acetyl lysine-dependent manner (Fig. 2d,e). Of note, YFV-C single lysine acetylation (Lys-4 or Lys-8) is sufficient for the BRD4-BD1 interaction. Furthermore, the interaction affinity of acetylated YFV-C histone mimic-containing peptide with BRD4 (362μM for YFV-C K4ac and 540μM for YFV-C K8ac) is in a similar range as the affinity of H4 K5ac or H4 K8ac peptides for BRD4 (Extended Data Fig. 5f).

The similarity between histone H4 and YFV-C N-terminal peptide binding to BD1 was further underscored by the analysis of high-resolution (1.42 to 1.53 Å) crystal structures of complexes of YFV-C containing K4acK8ac, K4ac and K8ac peptides to the BD1 bromodomain of BRD4 (Fig. 2f-h and x-ray statistics in Extended table S1). The three complexes showed high-resolution electron density for the bound YFV-C peptide backbone and side chains (Extended Data Fig. 7a-f).

The binding pocket analysis of YFV-C dual acetylated and YFV-C K4ac monoacetylated peptide show that the K4ac side chain inserts into the bromodomain binding pocket and is aligned in position through hydrogen bond formation with the side chain of asparagine 140 (N140), a feature common with earlier structures of acetyl lysine recognition by bromodomains^19,20^. Moreover, the peptide side chain of YFV-C arginine 3 (R3) hydrogen bonds with the side chain of aspartate 144 of BD1 in both complexes (Fig. 2f,g). The bound peptides are aligned in opposite directions in the YFV-C K4ac and K8ac complexes with BD1 (Extended Data Fig. 7g,h), resulting in the side chain of R3 in the YFV-C K8ac monoacetylated hydrogen bonding with the side chain of aspartate 145 in the BD1-YFV-C K8ac complex (Fig. 2h). Given that YFV-C R3 plays a key role in peptide-protein recognition, we determined the interaction affinity of BRD4 BD1 and acetylated YFV-C bearing arginine-to-alanine substitution at position 3 (R3A). We observed that R3A substitution in YFV-C peptides abolished the interaction of YFV-C to BRD4-BD1 regardless of lysine acetylation state (Extended Data Fig. 7i).

The ability of YFV-C H4-like motif (hereafter histone mimic) to interact with BRD4 as well as other BET proteins suggests a plausible mechanism of YFV-C-mediated interference with gene expression. Therefore, we compared gene expression changes associated with YFV-C and YFV-C K4RK8R. The gene expression profile of Huh-7, Huh-7 YFV-C and Huh-7 cells expressing YFV-C K4RK8R (Huh-7 K4RK8R) shows four gene clusters according to YFV-C histone mimic acetylation motif (histone mimic-dependent and independent) with around 40% of total changed genes being histone mimic-dependent (Fig. 3a). Induced histone mimic-dependent genes are enriched for extracellular matrix organization and cell morphogenesis while induced histone mimic-independent genes are enriched for autophagy, neutrophil activation and post-translational protein modification (Fig. 3b). Suppressed histone mimic-dependent genes are related to mRNA nonsense-mediated decay, protein targeting to ER and membrane (Fig. 3b), which are also among the most suppressed YFV-C processes (Fig.1g and Extended Data Fig. 2a). The expression profiles of Huh-7 YFV-C and Huh-7 YFV-C K4RK8R only share changes in the suppression of DNA replication and chromatin assembly processes, among the most changed biological processes (Extended Data Fig. 8a), indicating a key function of histone mimicry in biological processes modulation by YFV-C.

**Figure 3.**
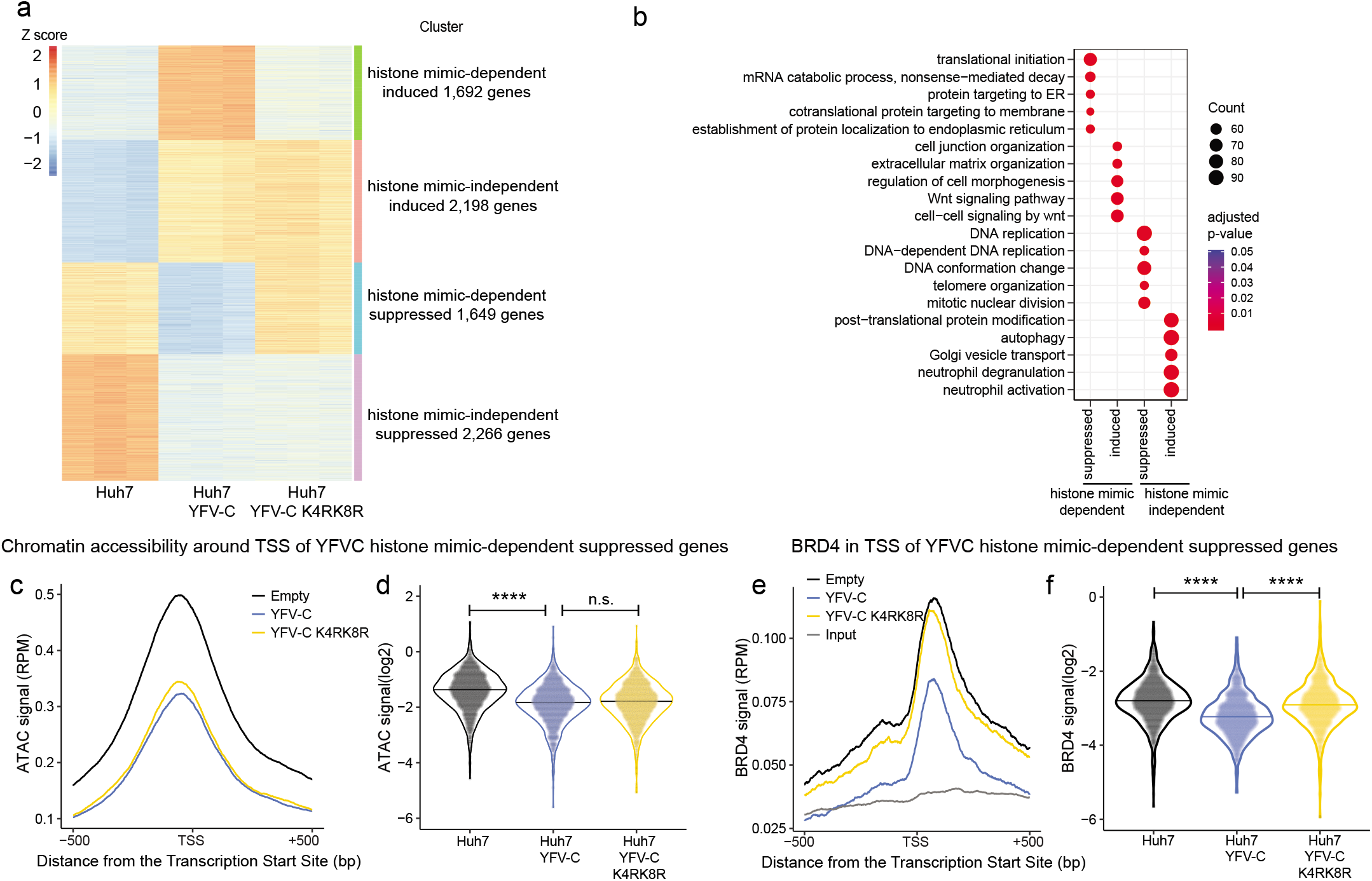
YFV-C modulates gene expression and BRD4 chromatin levels in a histone mimicdependent manner. **a,** Clustered heatmap of gene expression changes in Huh-7 cells, Huh-7 cells expressing YFV-C and Huh-7 cells expressing YFV-C with the K4RK8R mutations. The 4 clusters identified based on YFV-C and YFV-C K4RK8R changed genes were named histone mimicdependent suppressed or induced and histone mimic-independent suppressed or induced. **b,** Gene clusters were analyzed by gene enrichment on GO biological process terms. Circles sizes (Count) depict the number of genes in the gene cluster that overlap with GO terms. **c,** Genomic chromatin accessibility (ATAC signal) around the transcription start site (TSS) of 1,649 genes of histone mimic-dependent suppressed gene cluster for Huh-7, Huh-7 YFV-C and Huh-7 YFV-C K4RK8R were superimposed. **d**, Violin plot of the distribution of the ATAC signal of the distribution in **c**. Statistical differences were assessed by t test. **e,** Genomic chromatin levels of BRD4 measured by ChIP-seq (BRD4 signal) around the transcription start site (TSS) of 1,649 genes of histone mimicdependent suppressed gene cluster for Huh-7, Huh-7 YFV-C and Huh-7 YFV-C K4RK8R were superimposed. **f**, Violin plot of the distribution of the BRD4 signal of the distribution in **e**. Statistical differences were assessed by t test. Not significant (n.s) depicts p-value > 0.05 and **** depicts p-value < 10^-4^.

The capacity of YFV-C histone mimicry to modulate gene expression changes may be related to regulation of chromatin accessibility and/or BRD4 chromatin levels. Thus, using ATAC-seq, we compared chromatin accessibility of histone mimic-dependent genes upon YFV-C or YFV-C K4RK8R expression. No increased accessibility was observed on the TSS’s of induced histone mimic-dependent genes (Extended Data Fig. 8d,e), while a reduced chromatin accessibility was observed on TSS’s of suppressed histone mimic-dependent genes (Fig. 3c,d). Notably, YFV-C reduced chromatin accessibility regardless of histone mimicry. BRD4 genome-wide chromatin levels were quantified by chromatin immunoprecipitation sequencing (ChIP-seq) in Huh-7 cells and in Huh-7 cells expressing YFV-C or YFV-C K4RK8R. BRD4 levels at the TSS of genes that are up-regulated in a histone mimic-dependent fashion are not significantly changed upon YFV-C expression (Extended Data Fig. 8d-f). However, BRD4 levels at the TSS of genes that are down-regulated in a histone mimic-dependent fashion are significantly reduced upon YFV-C expression (Fig. 3e,f). Moreover, the impact of the wild-type YFV-C on BRD4 abundance at the TSS depends on the histone mimic as suggested by reduced BRD4 levels at the TSS compared to Huh-7 or Huh-7 YFV-C K4RK8R (Fig. 3e,f and Extended Data Fig. 8g). Accordingly, BRD4 chromatin levels achieve a higher positive correlation with gene expression than chromatin accessibility (Extended Data Fig.8h,I, Pearson r = 0.69 vs Pearson r= 0.48). Moreover, we addressed the link between BRD4 abundance at the individual gene loci and the presence of the histone mimic. We found that BRD4 binds to 5,165 gene loci in Huh-7 and/or Huh-7 cells expressing YFV-C with the majority (80%) localized at the TSS of genes (Extended Data Fig. 9a). Among the BRD4-associated loci, 660 loci (13%) showed changes in BRD4 levels in response to YFV-C with the majority of changes localized to the TSS (Extended Data Fig. 9b,c). Loci associated with BRD4 reduction are also associated with reduction in gene expression and loci associated with increased BRD4 are associated with increased gene expression (Extended Data Fig. 9d-f) in line with BRD4’s role in gene transcription. We estimate that ectopic expression of YFV-C yields 10^6^ molecules in the nucleus where approximately 1% can be found acetylated in either lysine 4 or 8 (Extended Data Fig. 4a). This amount of YFV-C could, potentially, suffice for targeting 25% of BRD4 molecules (based on the published data of 4 × 10^4^ BRD4 nuclear molecules in a related hepatocyte cell line)^21^.

As for gene expression changes, we observed both YFV-C histone mimic-dependent and - independent BRD4 changes. YFV-C histone mimic-associated reduction of BRD4 levels was observed on 131 loci (Fig. 4a). These loci may represent histone mimic-dependent YFV-C targets for BRD4 modulation and may encode transcriptional regulators that control the plethora of gene expression. To test that, we used the ChIP Enrichment Analysis (ChEA) database^22^ to probe for transcription factors associated with YFV-C histone mimic-dependent suppressed genes (Fig. 4b). Among the transcription factors identified, we observed that CREM and CCND1 have both reduced gene expression and BRD4 binding in a histone mimic-dependent manner (Fig. 4b,c).

**Figure 4.**
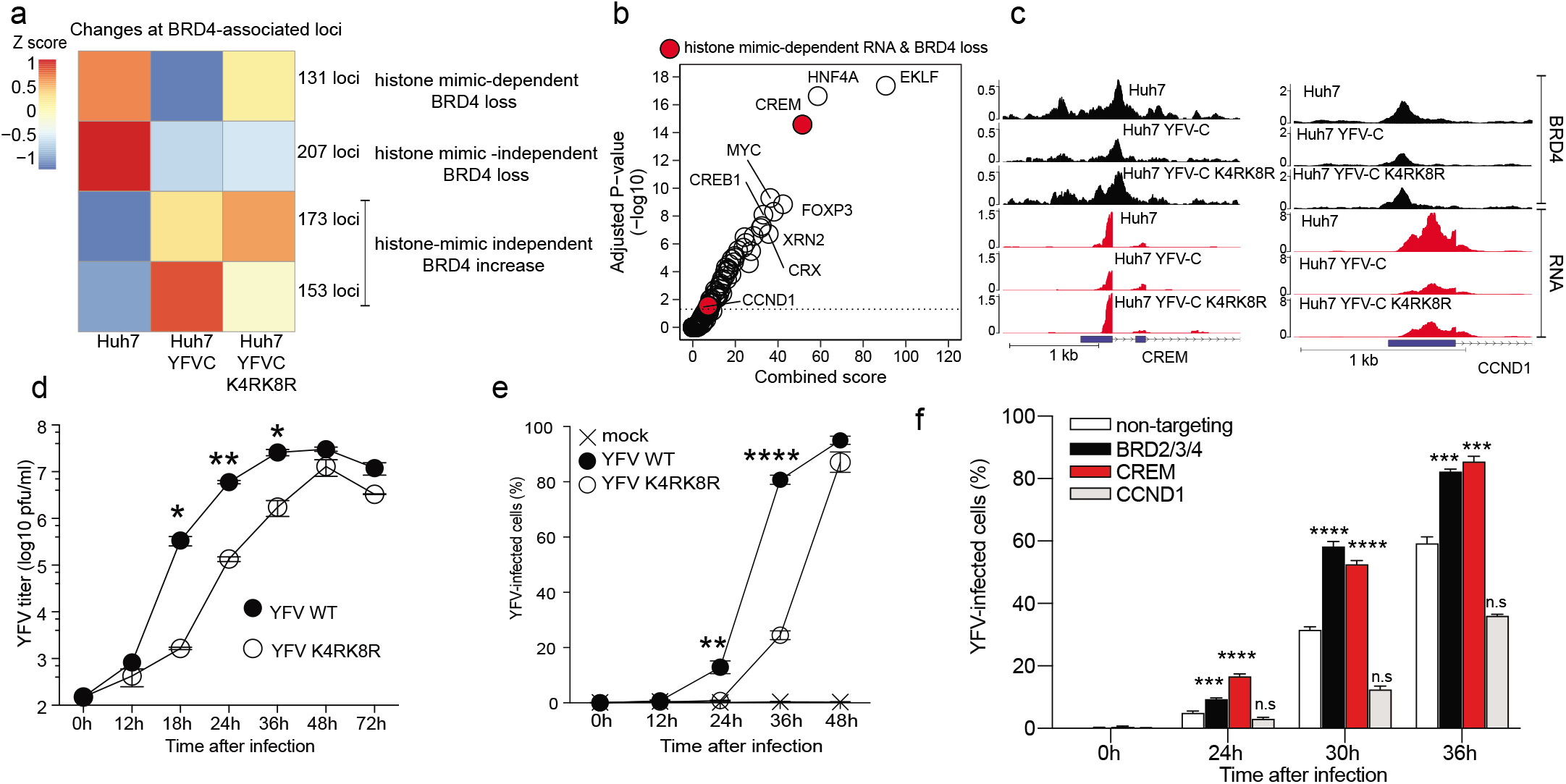
YFV-C can contribute to YFV replication via BET control of CREM expression. **a,** clustered average heatmap of BRD4 genomic loci level changes in Huh-7, Huh-7 YFV-C and Huh-7 YFV-C K4RK8R. **b,** ChEA 2016 Enrichr transcription factor binding analysis. Shown in red are transcription factors associated both with YFV-C histone mimic-dependent suppressed RNA and BRD4. **c**, Snapshot of BRD4 in genomic loci of the transcription factors shown to be associated both with YFV-C histone mimic-dependent suppressed RNA and BRD4. **d**, Huh-7 cells were infected (MOI=0.1) with YFV 17D WT or YFV 17D K4RK8R and YFV titers were determined in the culture supernatants, collected at the stated time, by plaque assay. Data is representative of 3 independent experiments. **e**, As in **d** but cells were collected at the indicated time, fixed, permeabilized and stained for YFV antigen prior to FACS analysis. Data is representative of 3 independent experiments. **f**, siRNA knockdown of CREM, CCND1 or triple knockdown of BRD2, 3 and 4 in Huh-7 cells was performed. Cells were infected (MOI 0.1) with YFV 17D 48h after siRNA transfection. Cells were collected at the stated time point, fixed, permeabilized and stained for YFV antigen prior to FACS analysis. Data is representative of 3 independent experiments. Statistical differences were assessed by t test. Not significantly increased (n.s) depicts p-value > 0.05 and **** depicts p-value < 10^-4^.

We hypothesized that the observed impact of the YFV-C histone mimic on gene expression, chromatin accessibility and BRD4 chromatin levels might be associated with changes in viral infectivity. We found that substitution of both lysine 4 and 8 to arginine in the YFV 17D strain (YFV K4RK8R), demonstrably reduces YFV infectivity as measured by plaque assay titers and virus spread measured by YFV intracellular staining (Fig. 4d,e). The reduced virus spread and titers observed in the YFV K4RK8R infected cells are not due to virus entry defects as *in vitro* transcribed viral RNA genome transfected into cells still yield reduced titers in the same frequency of infected cells (Extended Data Fig. 10a,b). We also confirmed that encapsulation of YFV K4RK8R was not affected by measuring the specific infectivity, the ratio of infectious virus particle to virus genome (Extended Data Fig. 10c). This result is in line with reports that around 40 N-terminal residues of YFV-C, which includes the histone mimic motif, is redundant for genome packaging and virus particle formation^23^. The YFV-C histone mimic does not overlap with the YFV genome cyclization region that plays a role in genome packaging and replication^24,25^. Additionally, YFV K4RK8R showed reduced plaque size in comparison to YFV WT (Extended Data Fig. 10d). Moreover, YFV K4RK8R showed bigger plaque sizes on later time points after infection (Extended Data Fig. 10e), suggesting a re-gained ability of the mutant YFV to recapitulate the YFV WT plaque size phenotype. To evaluate whether the increased plaque size observed on YFV K4RK8R was due to a genetic reversion of YFV-C K4RK8R mutations, we sequenced RNA from virus particles released into the culture media. We observed that the YFV-C sequence showed evidence for genetic reversion of arginine-to-lysine at positions 4 and 8, suggesting a selective pressure for the maintenance of lysines at position 4 and 8 of YFV-C (Extended Data Fig. 10f). Indeed, this can be observed by the increased conservation of this segment among different YFV strains (Extended Data Fig. 3e). The crystal structure of the YFV-C peptide complexed with BRD4 points to arginine 3 of YFV-C as a virus-specific determinant of the histone mimic interaction with BET (Fig. 2f-h). The infectivity of YFV R3A is greatly reduced in comparison to YFV (Extended Data Fig. 10g). Importantly, YFV replication attenuation by mutations in the YFV-C-BRD4 interaction surface (K4RK8R and R3A) is conserved in *Aedes aegypti* mosquito cells, an obligate host through which YFV cycles (Extended Data Fig. 10h). In agreement with YFV-C histone mimic control of CREM expression, knockdown of CREM and BET (BRD2, 3 and 4), but not CCND1, increased YFV replication (Fig. 4f) as well as pharmacological inhibition of BET proteins (Extended Data Fig. 10i).

Analysis of the biological processes suppressed by YFV-C and identified as histone mimic-dependent, such as “mRNA catabolic process – nonsense-mediated decay” and others, show 25–30% of overlap by YFV-C in histone mimic-dependent and CREM-bound genes (based on published data extracted by ChEA) (Extended Data Fig. 11a-e). Other processes suppressed by YFV-C and identified as histone mimic-independent show little overlap with CREM-bound genes (Extended Data Fig. 11f-i).

Collectively our data point to a role for YFV-C in impairment of BRD4-induced expression of CREM, that in turn regulates different genes in biological processes that impact YFV replication in Huh-7 cells. Of note, nonsense-mediated decay (NMD) has been implicated in the control of flavivirus replication^10–12^. Identification of YFV-C arginine 3 as an important determinant of YFV virulence may also enable development of novel antiviral therapeutics that will inhibit YFV-C, but not cognate histone, interaction with BET. Our data suggest that a significant and infection-relevant effect of YFV-C on gene expression is carried by the N-terminal portion of YFV-C that is homologous to histone H4. The similarity between the YFV-C and histone H4 also suggests that YFV-C can function as a viral equivalent of synthetic BET inhibitors, such as I-BET or JQ1, that have a well-known capacity to interfere with signal induced gene expression, including expression of type I interferon^26^. Therefore, identification of histone “readers” that can bind to viral sequences may inform development of novel selective epigenetic drugs that target viral infection and inflammation.

---

**Extended Table S1.**
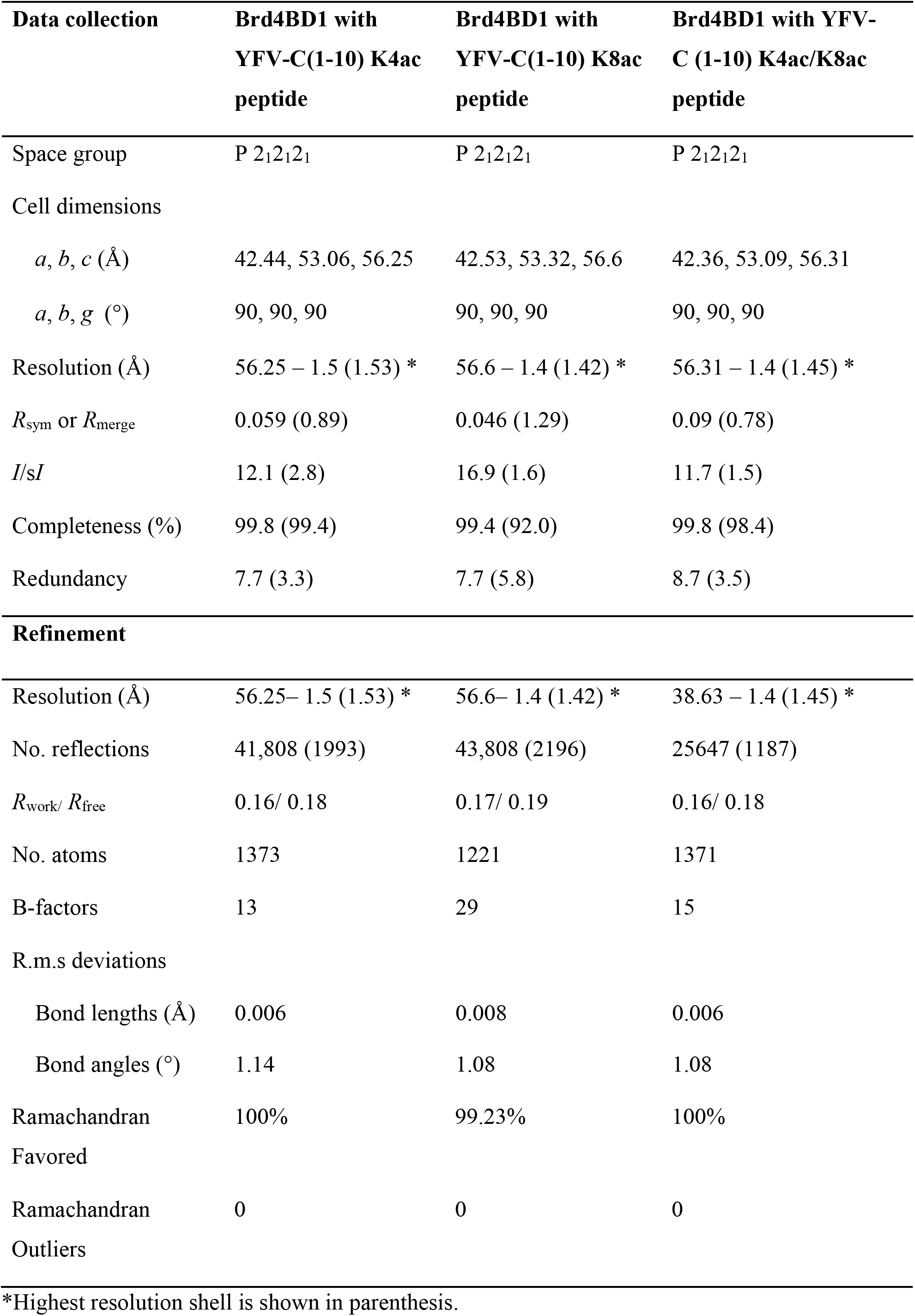
X-ray data collection and refinement statistics.

**Extended figure 1.**
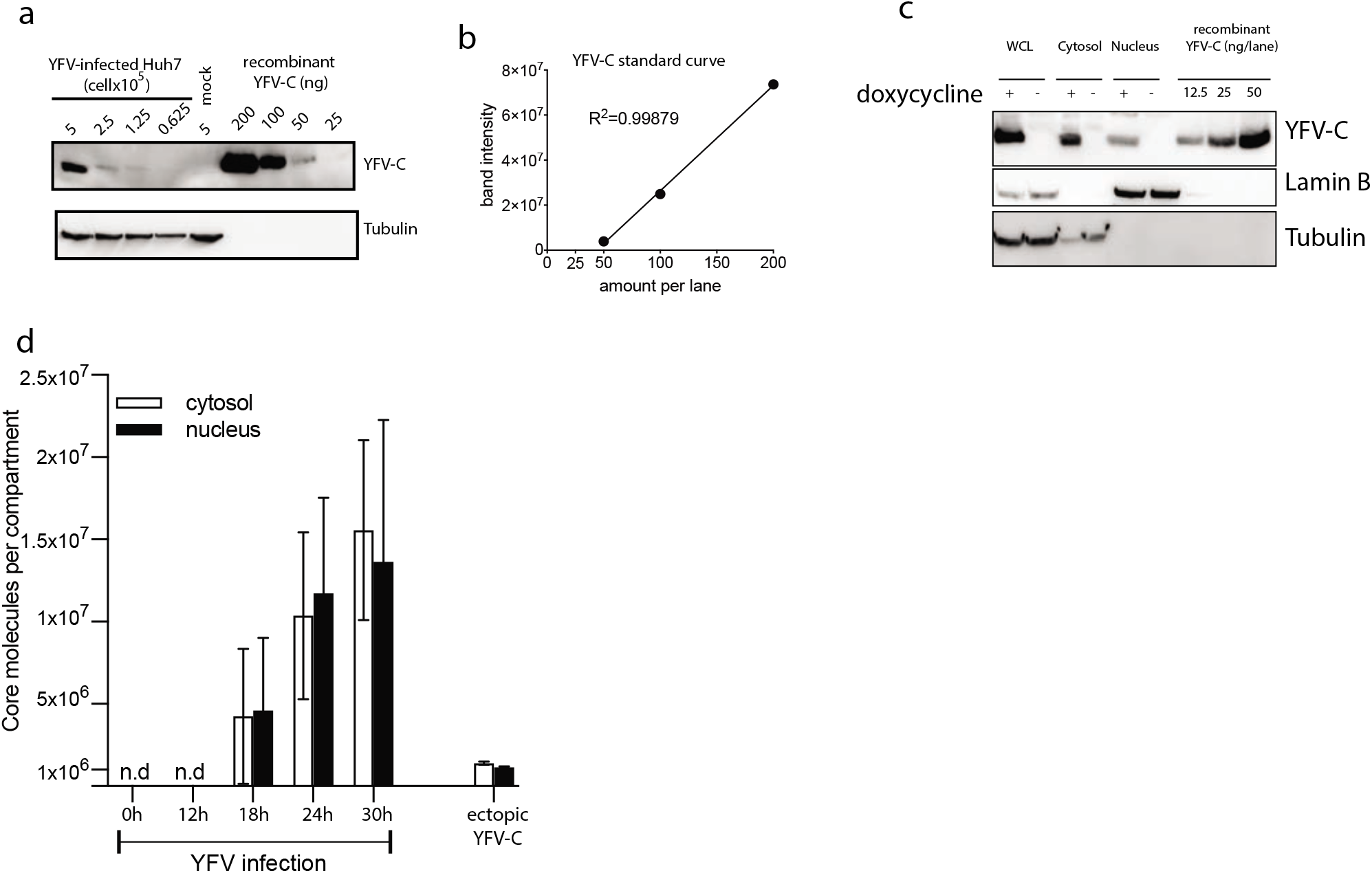
Quantification of YFV-C in Huh-7-infected and ectopic-expressing cells. **a**, Western blot analysis of YFV-C amounts in whole cell lysates from different numbers of Huh-7 cells infected with YFV 17D MOI 10 for 24 hours (left panel) and the standard curve derived from the western blot with recombinant purified YFV-C protein (right panel). Representative of 3 independent experiments. **b**, Example of a standard curve of YFV-C titration by Western blot. **c**, Western blot analysis of YFV-C amounts levels in different cellular compartments of Huh-7 ectopically and inducibly expressing YFV-C untreated or treated with doxycycline 10μg/ml for 24 hours. **d**, Quantification of the number of YFV-C molecules in the cytosol and nuclei of infected cells where a mass of 1ng of YFV-C (11.49 kDa) equals 5.23 × 10^10^ molecules. Error bars represent standard deviation of 3 independent experiments.

**Extended figure 2.**
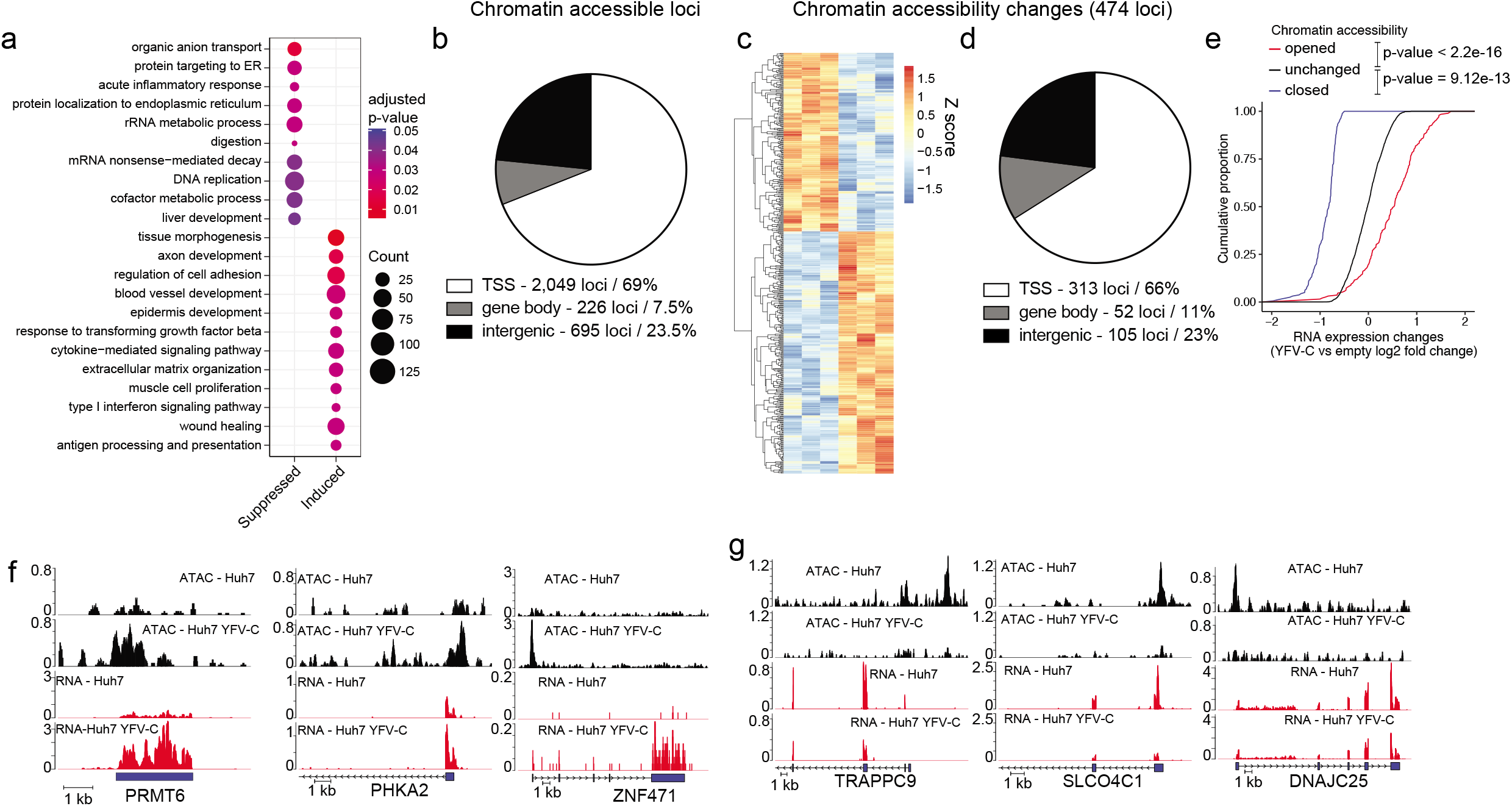
YFV-C-mediated gene expression and chromatin accessibility changes correlation. **a**, Plot of the 10 most changed GO biological processes terms (by adjusted p value) found to be suppressed or induced by YFV-C ectopic expression. The circle sizes correspond to the number of YFV-C affected genes in the term. **b,** Chromatin accessible sites were assessed by its genomic location; transcription start site (TSS), gene body or intergenic regions. **c,** Heatmap of all chromatin accessibility changes. **d,** Chromatin accessibility changes were assessed by their genomic locations; transcription start site (TSS), gene body or intergenic regions. **e,** Genes with increased (open), decreased (closed) or unchanged chromatin accessibility (assessed by adjusted p value < 0.05) were plotted by its cumulative distribution of RNA expression changes. **f,** Example tracks at chromatin gene levels of chromatin accessibility and RNA levels changes for YFV-C-induced or **g,** decreased RNA and chromatin accessibility (assessed by adjusted p value < 0.05).

**Extended figure 3.**
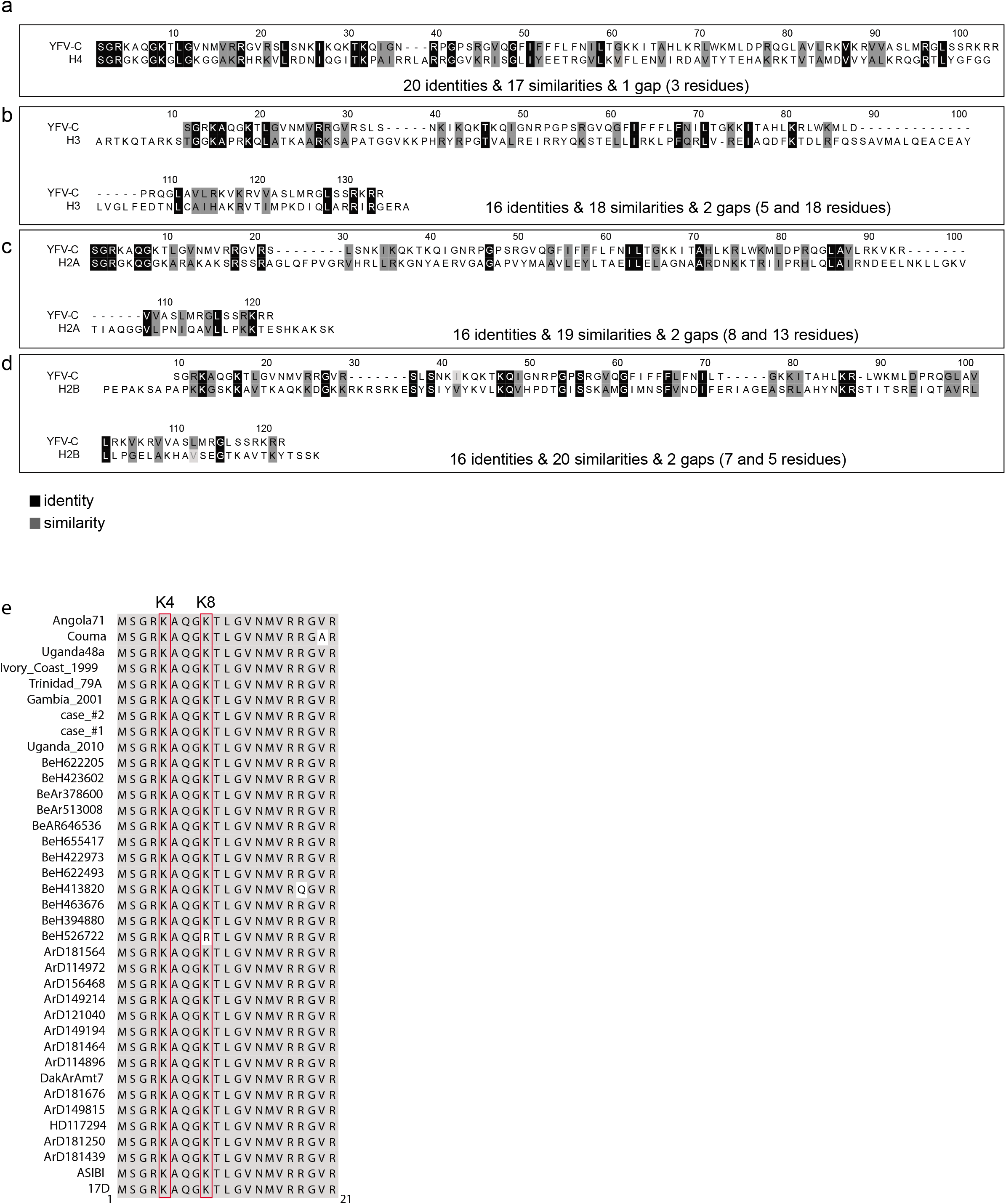
YFV-C resembles histones and has a conserved H4-like motif across strains. Amino acid alignment using ClustalW of YFV-C (accession NP_775998) and human histone **a**, H4 (accession NP_778224) **b**, H3.1 (accession NP_003522) **c**, H2A type 2-C (accession NP_003508) and **d**, H2B type 2-E(accession NP_003519). **e**, The 20 N-terminal amino acids (excluding the first methionine) of several different strains of YFV obtained on the Virus Pathogen Resource (ViPR) database were aligned by ClustalW.

**Extended figure 4.**
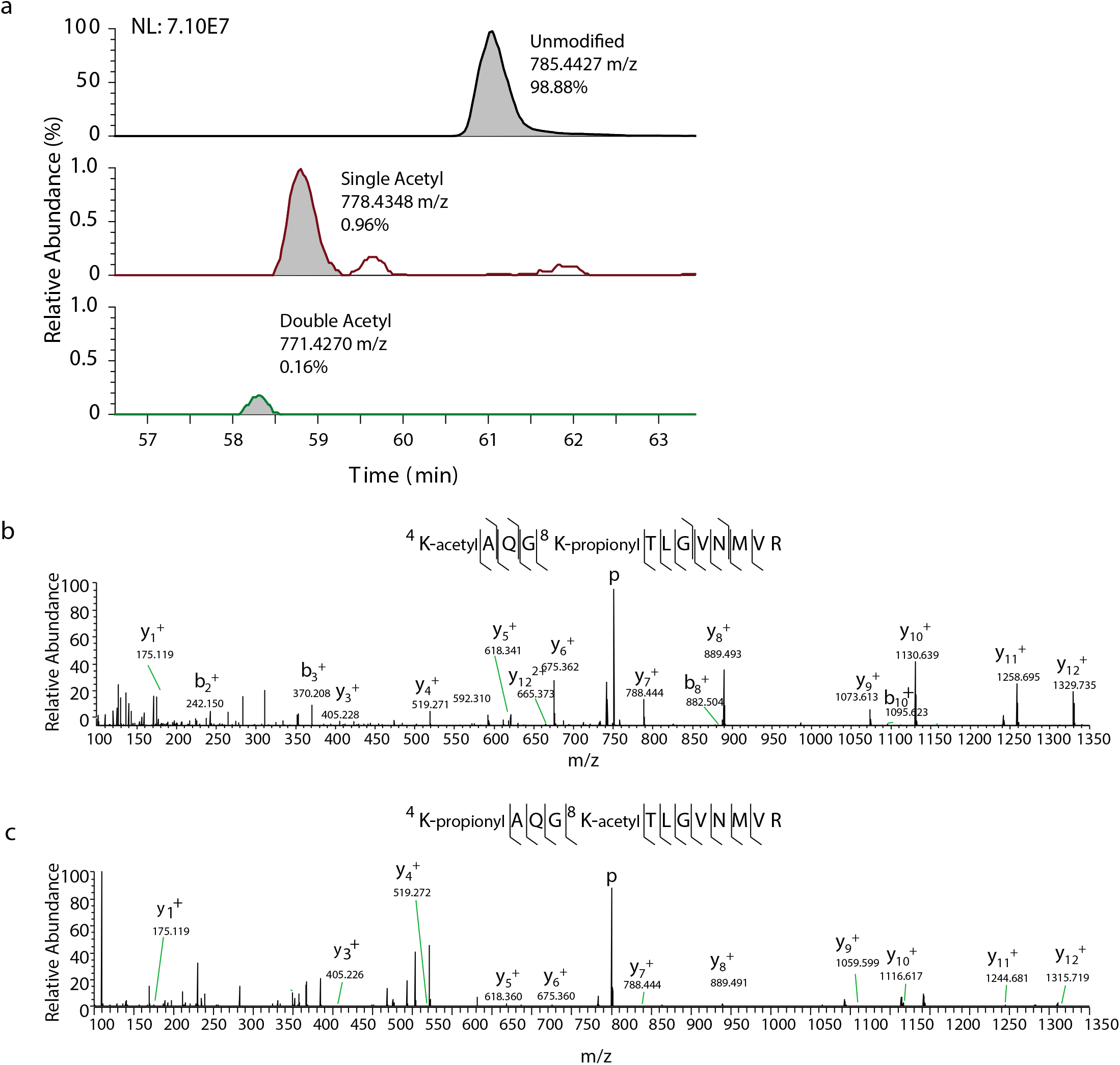
Mass spectrometry analysis of lysine acetylated YFV-C. **a**, extracted ion chromatograms of the unmodified and singly acetylated form of tryptic peptide KAQGKTLGVNMVR from the YFV-C protein. The relative abundance based on area under the curve of the base peak that corresponded to mass/charge 785.4427 and 778.4348 at elution times shown in the graph was used to calculate a relative abundance of 98.88% for unmodified peptide and about 0.96% for singly acetylated (either at K4 or K8). This is reflected in a 2-log difference in the normalized intensity shown at the upper left of the graph. A peak with m/z corresponding to the diacetylated form was also detected but could not be validated at MS/MS level due to interference from product ions of a 770 m/z species. **b**, T-Rex Flp-In HeLa cells carrying YFV-C-3xFlag were treated for 24 hours with 1μg/ml of doxycycline prior to nuclei isolation, Flag immunoprecipitation and polyacrylamide gel separation. Excised band was analyzed by MS/MS. Non-acetylated lysines were chemically modified by propionylation to protect from cleavage during trypsin digestion. Fragment b and y ions are marked. Tandem MS spectra of the tryptic peptide of YFV-C doubly charged Lysine 4 and **c**, Lysine 8 acetylated peptide: KAQGKTLGVNMVR. Non acetylated lysines are propionylated. Fragment ions (y and b) are marked. Based on signals the K4 acetylation form is predominant.

**Extended figure 5.**
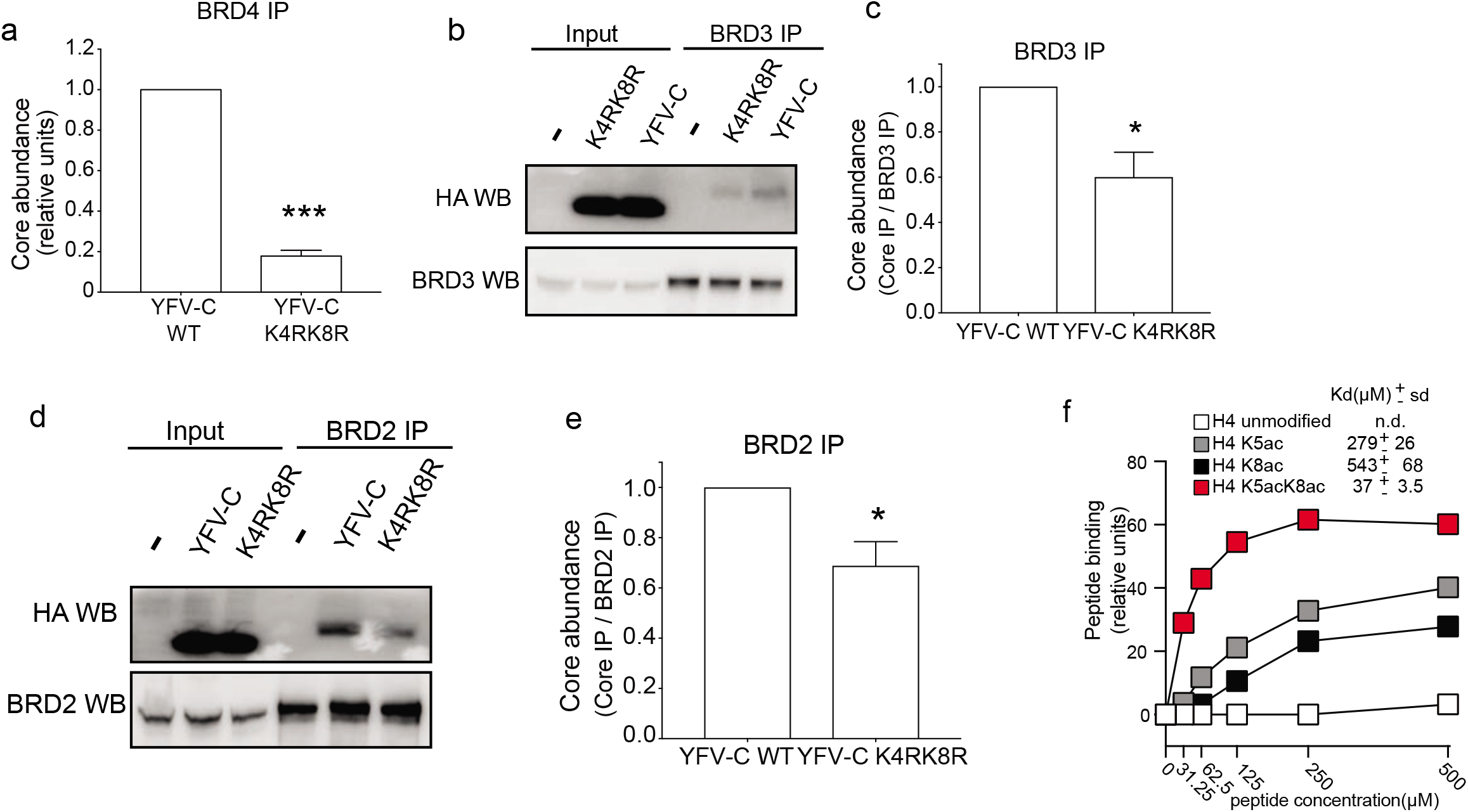
YFV-C binds to BET in a lysine 4- and 8-dependent manner. **a**, Densitometry analysis quantification of the YFV-C -Flag and BRD4 bands shown in Fig. 2c using the ImageLab Biorad software. Data is average of 3 independent experiments. Error bars represent standard error of the mean. **b**, T-Rex Flp-In HeLa cells carrying an empty vector (empty), vector expressing 3XFlag epitope-tagged YFV-C (WT) or 3XFlag tagged YFV-C with K4 to R and K8 to R mutations (K4RK8R) were treated with 1μg/ml of doxycycline for 24 hours for YFV-C gene expression induction prior to cell lysis and BRD3 immunoprecipitation. **c**, As in **a** but with quantification of the YFV-C -Flag and BRD3 bands shown in **b**. **d**, As in b, but with BRD2 immunoprecipitation. **e**, As in **a** but with quantification of the YFV-C -Flag and BRD3 bands shown in **d**. **f**, Surface plasmon resonance (SPR) equilibrium plots of interactions of lysine acetylated histone H4 (residues 1-11) with bromodomain 1 of BRD4. The constant of dissociation (Kd) was calculated based on the Langmuir fit of the data. Data is representative of at least 3 independent experiments and error bars represent standard deviation.

**Extended figure 6.**
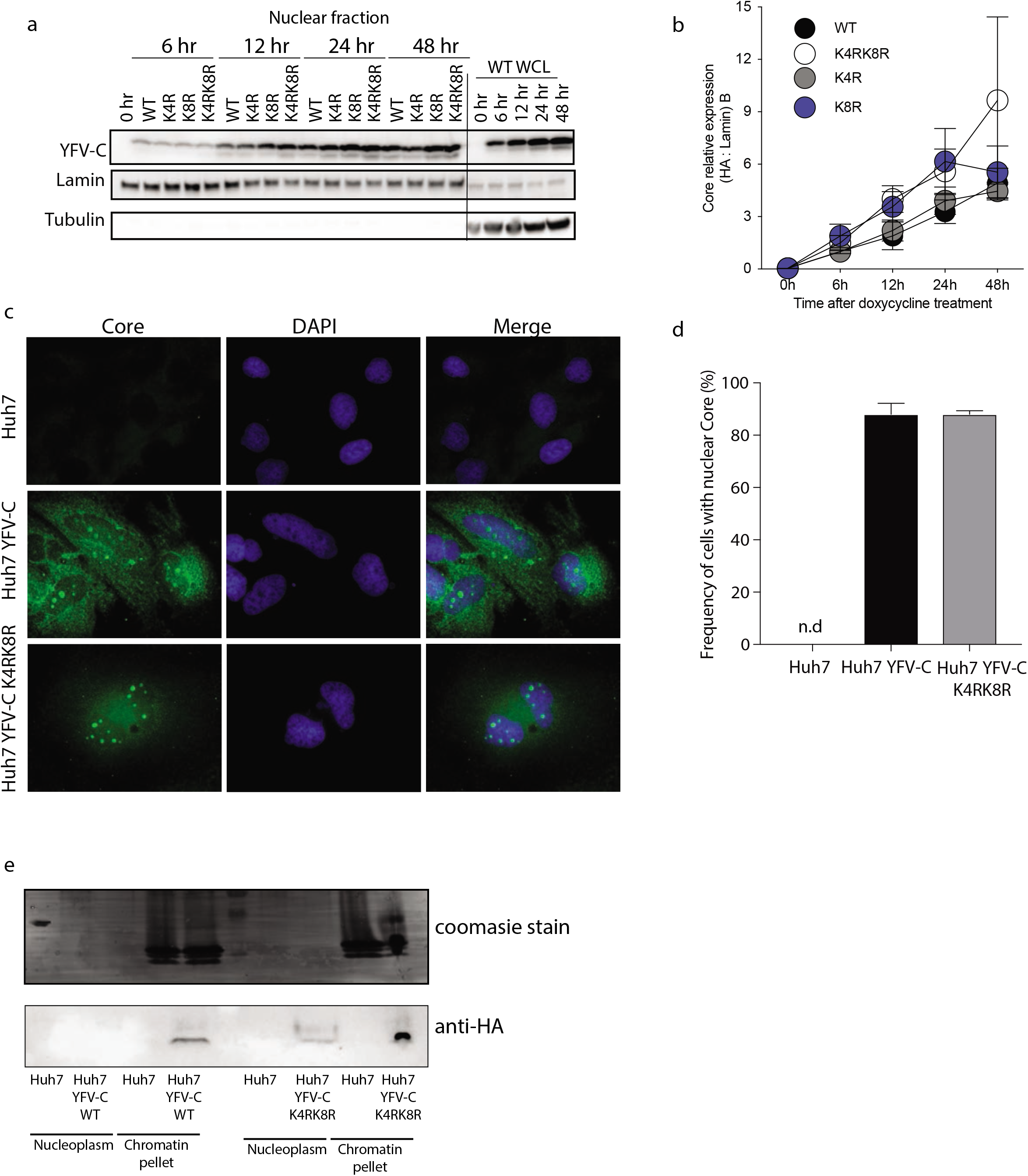
YFV-C lysine 4 and 8 acetylation do not alter subcellular localization. **a**, T-Rex Flp-In HeLa cells carrying an empty vector (empty), HA epitope-tagged YFV-C (WT), HA-tagged YFV-C with K4 to R mutation (K4R), K8 to R mutation (K8R) or K4 to R and K8 to R mutations (K4RK8R) were treated with 1μg/ml of doxycycline for the stated time prior to cell fractionation. HA expression, lamin B and a-tubulin was assessed by western blot. Data is representative of 3 independent experiments. **b**, HA expression levels were quantified in relation to lamin B levels. Data is representative of 3 independent experiments. **c**, Huh-7, Huh-7 YFV-C and Huh7 YFV-C K4RK8R were treated with 10μg/ml of doxycycline for 24 hours prior to fixation and immunofluorescence staining with rabbit anti-YFV-C and DAPI. Cells were analyzed by confocal microscopy and **d**, the frequency of nuclear YFV-C (green) positive was quantified. There were no positive cells on Huh-7 samples. Data is representative of 3 independent experiments and error bars represent standard error of the mean. **e**, NUN fractionation of nuclei into chromatin and nucleoplasm fractions upon ectopic expression of YFV-C or YFV-C K4RK8R in Huh-7 cells. Fractions were analyzed by coomassie staining of the gel (top) and western blot using anti-HA antibody (bottom).

**Extended figure 7.**
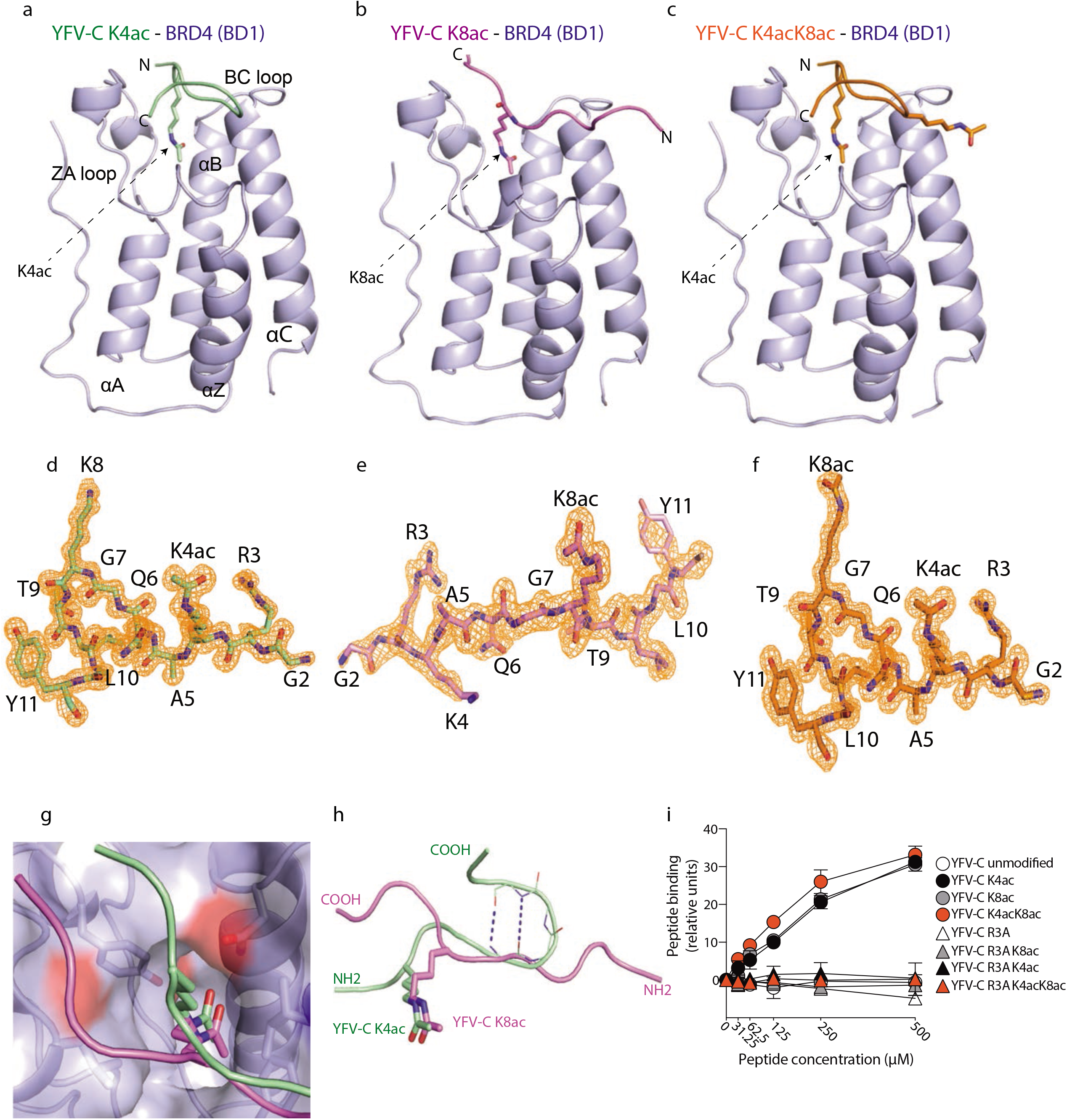
X-ray crystal structures of BRD4-BD1 and YFV-C lysine-acetylated peptide complexes. **a – c**, X-ray structures of YFV-C (1-11) peptides containing K4ac (panel a), K8ac (panel b) and K4ac/K8ac (panel c) peptides bound to the pocket of the BD1 bromodomain of Brd4. **d – f**, Tracing of the bound BD1-bound YFV-C (1-11) Kac-containing peptides in 2Fo-Fc omit maps (1.2 σ) in their BD1 complexes. The data are shown for bound K4ac (panel d), K8ac (panel e) and K4ac/K8ac (panel f) peptides in their BD1 complexes. **g,** Superposing the YFV-C (1-11) K4ac (in green) and K8ac (in magenta) peptide complexes bound to the BD1 bromodomain of Brd4, with positioning of YFV-C (1-11) K4ac and K8ac in the same binding pocket of the bromodomain of BD1. **h,** Opposite directionalities of the peptide chains of YFV-C (1-11) containing K4ac and K8ac in their complexes with the bromodomain of BD1. **i,** Surface plasmon resonance (SPR) equilibrium plots of interactions of lysine acetylated YFV-C with bromodomain 1 of BRD4. The constant of dissociation (Kd) was calculated based on the equilibrium between 45 and 50 seconds. Data is representative of at least 3 independent experiments and error bars represent standard deviation.

**Extended figure 8.**
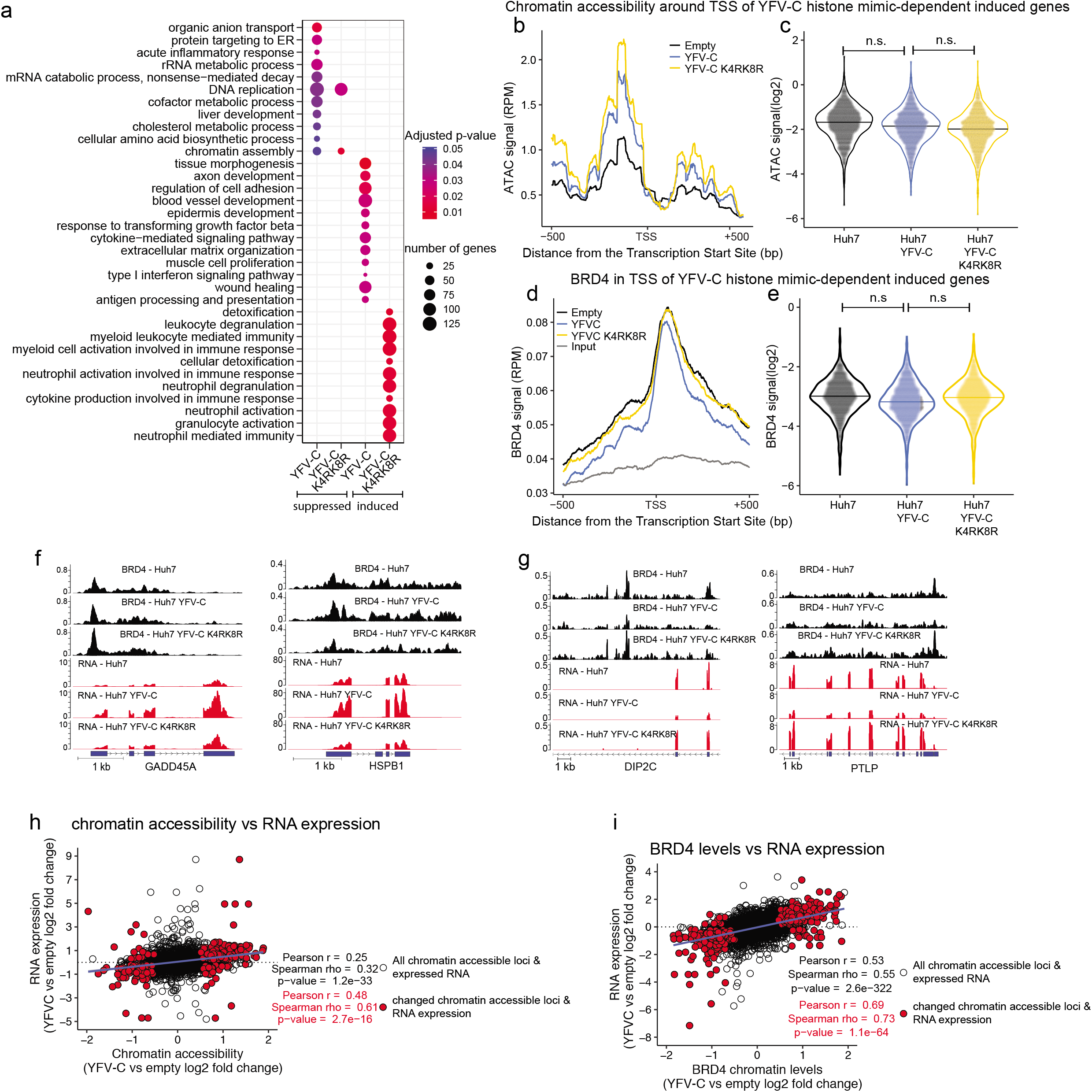
BRD4 and chromatin accessibility correlation to gene expression changes induced by YFV-C. **a,** All expressed RNA in Huh-7 cells expressing YFV-C or YFV-C K4RK8R was ordered by rank of gene expression differences to Huh-7 cells. The ranked list was analyzed by GSEA for suppressed or induced GO biological process terms. **b,** Genomic chromatin accessibility (ATAC signal) around the transcription start site (TSS) of 1,692 genes of histone mimic-dependent induced gene clusters for Huh-7, Huh-7 YFV-C and Huh-7 YFV-C K4RK8R cells were superimposed. **c**, Violin plot of the distribution of the ATAC signal of the distribution in **b**. Statistical differences were assessed by t test. **d,** Genomic chromatin levels of BRD4 measured by ChIP-seq (BRD4 signal) around the transcription start site (TSS) of 1,649 genes of histone mimic-dependent suppressed gene cluster for Huh-7, Huh-7 YFV-C and Huh-7 YFV-C K4RK8R cells were superimposed. **e**, Violin plot of the distribution of the BRD4 signal of the distribution in **d**. Statistical differences were assessed by t test. n.s depicts not significant. **f,** Genomic tracks of BRD4 and RNA levels changes for YFV-C-induced or **g,** decreased RNA expression (adjusted p-value < 0.001). **h**, Dot plot of chromatin accessibility changes versus RNA expression. Red dots represent genes with statistical differences between Huh-7 YFV-C and Huh-7 cells in both chromatin accessibility and RNA expression. Black dots represent all genes expressed with corresponding chromatin accessibility. Linear regression line is in blue with Pearson and Spearman statistical correlation written in the graph for total (black) or changed (red) genes. **i**, as h but showing dot plot and correlation of BRD4 chromatin levels and RNA expression.

**Extended figure 9.**
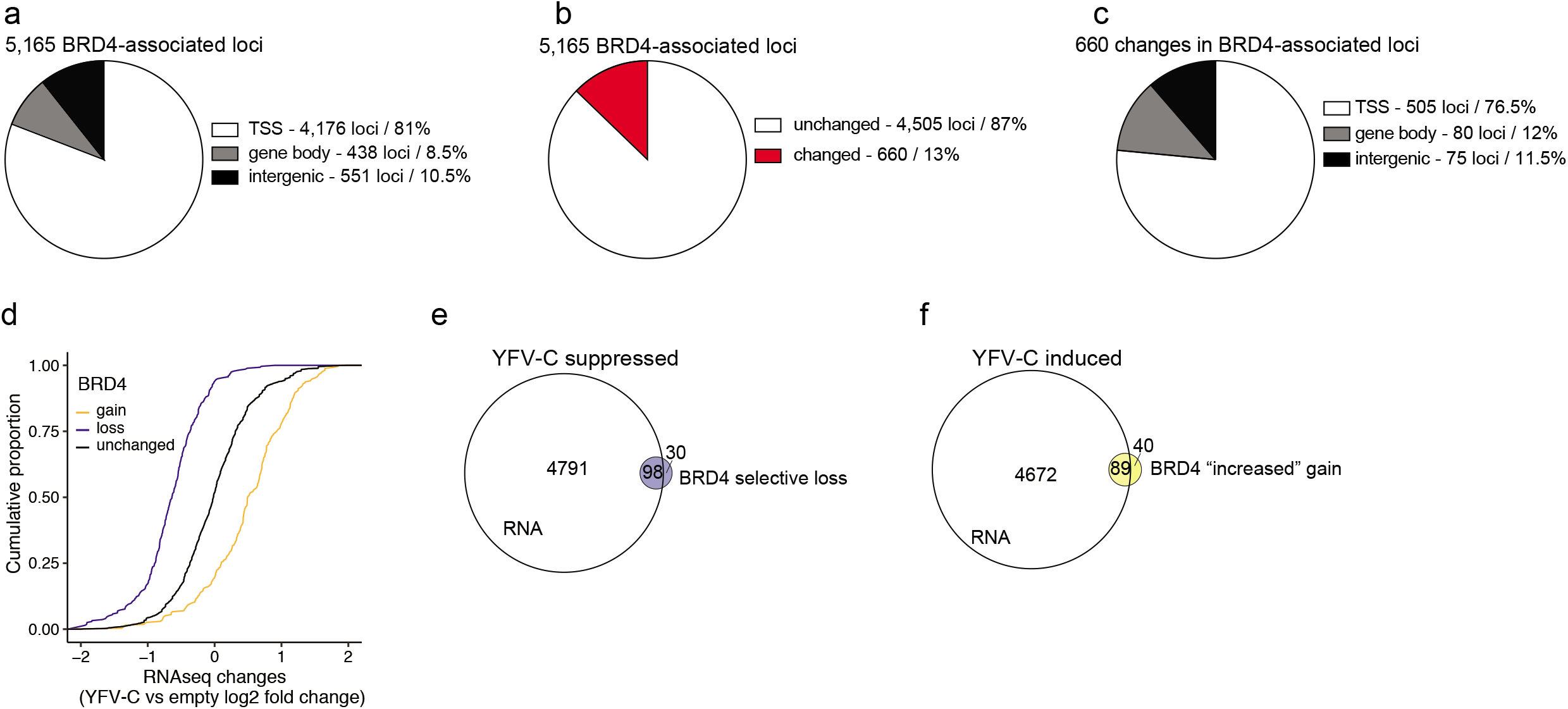
YFV-C modulation of BRD4 genomic levels correlates with modulation of gene expression. **a**, Genomic distribution of all the BRD4-associated loci observed either in Huh-7 or Huh-7 YFV-C -expressing cells. **b**, Genomic distribution of BRD4-associated loci changed upon 48 hours of YFV-C expression in Huh-7 cells. **c**, Genomic distribution of changed BRD4-associated loci observed either in Huh-7 or Huh-7 YFV-C-expressing cells. **d,** cumulative distribution of the fold change in BRD4 levels (Huh-7 YFV-C vs Huh-7) on genes associated with increased or decreased RNA expression. Statistical differences were assessed by Kolmogorov-Smirnov test comparing all genes with genes associated with increased or decreased BRD4 peaks. **e**, Venn diagram of genes associated with diminished BRD4 levels and YFV-C suppressed and **f**, genes associated with increased BRD4 levels and YFV-C induced.

**Extended figure 10.**
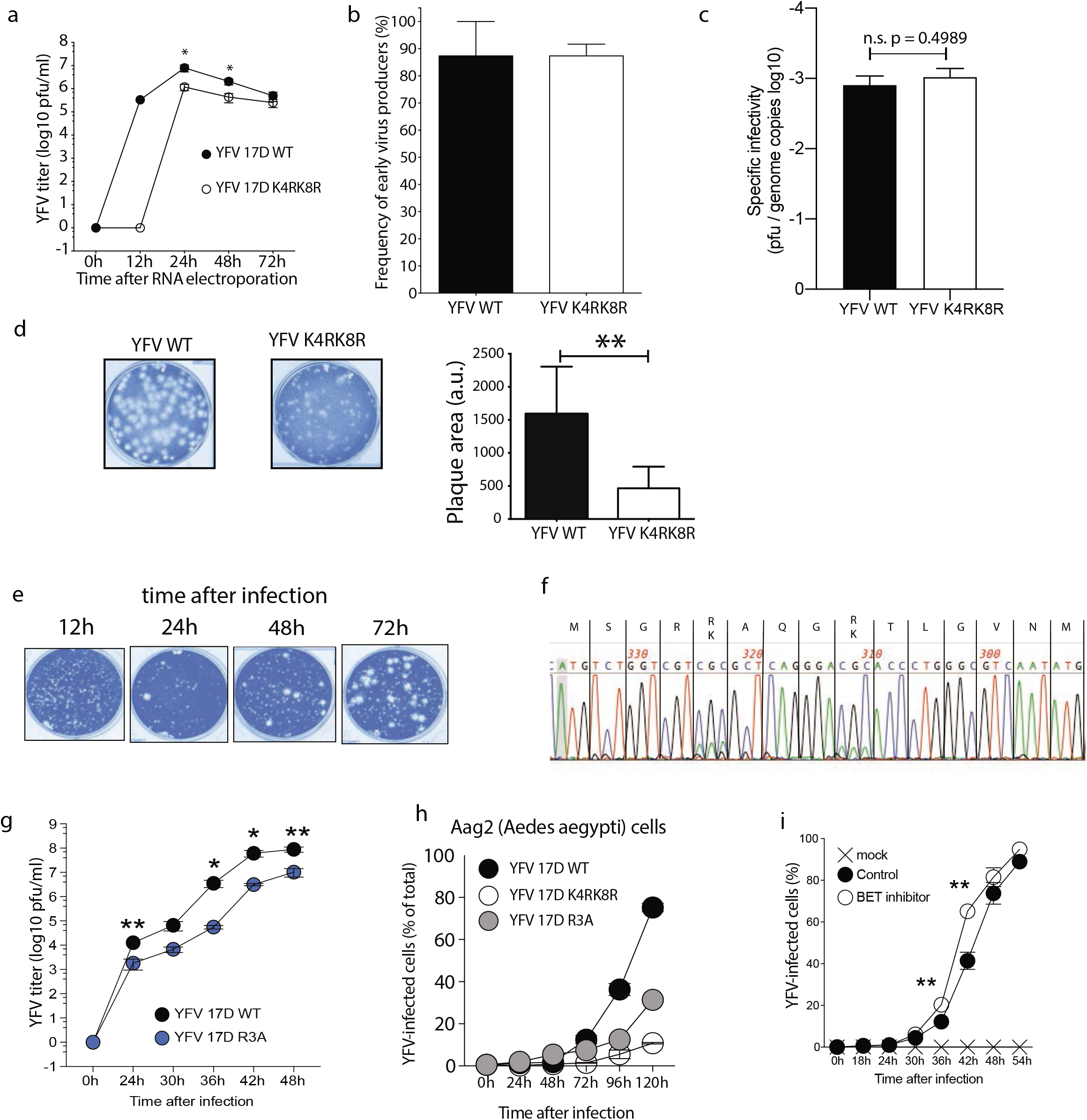
YFV 17D mutant virus attenuation is conserved in mosquito cells and is not due to virus entry or encapsidation defects. **a**, BHK-J cells (8 × 10^6^ cells) were electroporated with 5 μg of *in vitro* transcribed RNA derived from AflII linearized YFV 17D WT or YFV 17D K4RK8R infectious clone DNA and YFV titers were determined in the supernatants of culture, collected at the stated time, by plaque assay. Data is representative of 2 independent experiments. **b**, Different numbers of electroporated cells from **a** were plated with non-electroporated cells for 6 hours prior to agarose layering. The number of electroporated plated cells divided by the number of observed plaques is the frequency of early virus producers. Data is representative of 3 independent experiments. **c**, Huh-7 cells were infected (MOI=0.1) with YFV 17D WT, YFV 17D K4RK8R and YFV 17D R3A and YFV titers were determined in the culture supernatants collected at the stated time, by plaque assay and by quantitative PCR using YFV NS3 primers and PCR product for absolute quantification of YFV genome copies. The specific infectivity was obtained by dividing the plaque forming units of the plaque assay by the amount of virus genome copies present in the supernatant. Data is average of 2 independent experiments. Statistical differences were assessed by unpaired t-test. **d**, The size of the plaques formed by YFV 17D WT and YFV 17D K4RK8R, as in **b**, was measured by ImageJ software. **e**, Examples of different sizes of the plaques of YFV 17D K4RK8R obtained from different times of Huh-7 infected (MOI=0.1) cultures. **f**, RNA was isolated from supernatant collected 24 hours after YFV 17D K4RK8R infection using QIAamp viral RNA mini kit and reverse transcription PCR was performed. The YFV genome was amplified using YFV-C primers and sequenced after gel extraction. **g**, Huh-7 cells were infected with YFV 17D WT or YFV 17D R3A (MOI=0.1). YFV titers were determined in the supernatants of the culture, collected at the stated time, by plaque assay. Data is representative of 3 independent experiments. **h**, Aedes aegypti Aag2 mosquito cell line was infected (MOI=1) with YFV 17D WT, YFV 17D K4RK8R and YFV 17D R3A. Cells were collected at the indicated times, fixed, permeabilized and stained for YFV antigen prior to FACS analysis. Data is representative of 3 independent experiments. **i,** Huh-7 cells treated with 0.3μM JQ1 for 1 hour were infected as in **c** with YFV 17D and the treatment with JQ1 was continued throughout the course of infection. Cells were collected at the indicated times, fixed,permeabilized and stained for YFV antigen prior to FACS analysis. Data is representative of 3 independent experiments. Statistical differences were assessed by t-test. ** p-value < 0.01.

**Extended figure 11.**
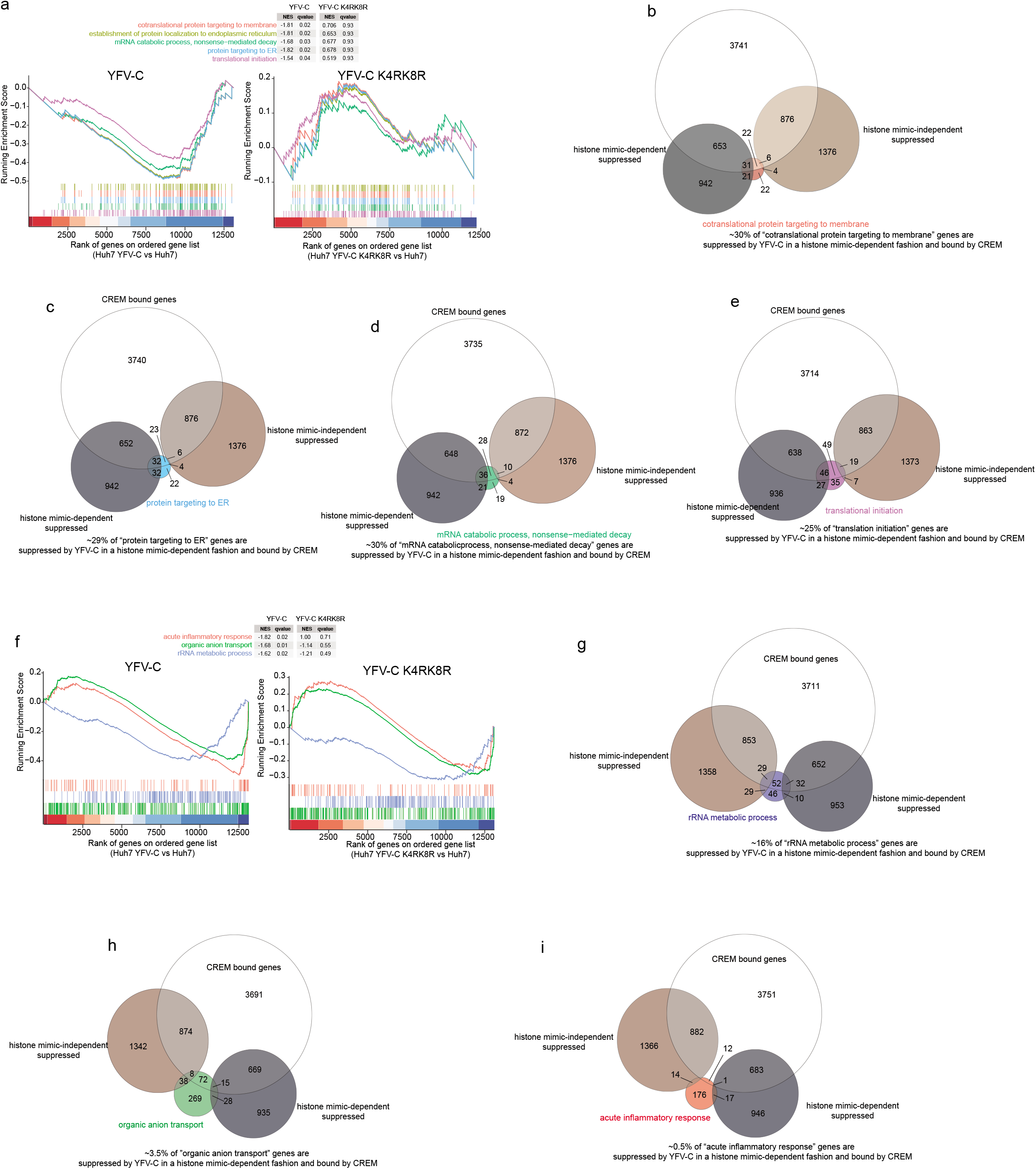
CREM involvement on YFV-C-modulated biological processes. **a**, GSEA plots of representative biological processes modulated by YFV-C in a histone mimic-dependent manner that show more than 25% of overlap of CREM-bound genes and suppressed YFV-C histone mimic-dependent genes. **b**, Venn diagram of CREM-bound genes”, modulated by YFV-C in a histone mimic-dependent or independent fashion and involved in “cotranslational protein targeting to membrane”, **c**, “protein targeting to ER”, **d**, “mRNA catabolic process, nonsense-mediated decay”, **e**, “translation initiation”. **f**, GSEA plots of representative biological processes modulated by YFV-C in a histone mimic-dependent manner that show less than 20% overlap of CREM-bound genes and suppressed YFV-C histone mimic-dependent genes. **g**, Venn diagram of CREM-bound genes”, modulated by YFV-C in a histone mimic-dependent or independent fashion and involved in “rRNA metabolic process”, **h**, “organic anion transport”, **i,** “acute inflammatory response”.

## Methods

### Cells

Huh-7, Huh-7 .5 and BHK-21 cells were cultured in Dulbecco’s minimal essential medium (DMEM) (Gibco /Thermo Fisher Scientific) supplemented with 2mM L-glutamine (Gibco /Thermo Fisher Scientific), non-essential amino acids (Gibco /Thermo Fisher Scientific), penicillin-streptomycin (Gibco /Thermo Fisher Scientific) and 10% fetal bovine serum (Hyclone). The T-Rex HeLa cell line (Thermo Fisher Scientific) was cultured in DMEM, L-glutamine, penicillin-streptomycin, non-essential amino acids and 10% Tetracycline-free FBS (Clonetech) and HeLa Trex cell line expressing YFV-C was generated according to the manufacturer’s instructions with pcDNA5/FRT/TO and pOG44 (Thermo Fisher Scientific). Huh-7 Tet on cells were generated by transduction of Huh-7 cells with lentivirus generated using the pCW57.1 plasmid (gift from David Root, Addgene plasmid # 41393).

### Virus production and viral assays

The YFV 17D K4RK8R and R3A were generated on the pACNR-FLYF17D-Da, a derivative of pACNR-FLYF-17D^1^ modified to use an AflII site for production of run-off RNA transcripts. Site-directed mutagenesis was conducted using a QuikChange Mutagenesis Kit (Stratagene) and virus stocks were generated with the same protocol as the wild-type virus. YFV 17D stocks were made by electroporation of 6 ×10^6^ to 8 ×10^6^ BHK-J cells with in vitro-transcribed RNA (5 μg) generated from AflII-linearized the pACNR-FLYF17D-Da as described previously^1,2^. The titers of the stocks were determined on BHK-J and Huh-7 cells after serial dilution as previously described^3^. Plaques were enumerated by crystal violet staining after 72 h. For YFV infections, the MOI was based on the titers obtained on Huh-7 cells. For the infectious center assay^4^, a sample of the electroporated cells was first diluted 1:100 and 1ml was added to a BHK-J cell monolayer (5 x10^5^ cells/well, 6 well plate) and then further diluted 1:10, and 2, 20, 200, and 1ml each was added to a BHK-J cell monolayer. After 4 h of incubation, the cells were overlaid with 0.6% agarose in MEM containing 2% FBS and incubated at 37°C. After 72h, infectious centers were enumerated by crystal violet staining as described above.

### Virus infection

Huh-7 cells were incubated with the corresponding MOI with 150μL/well in a 12-well format and 200μL/well in the 6-well plate format. Plates were rocked every 12 minutes for 1 hour and virus was discarded, and fresh media was added to wells.

For western blot analyses 4-5 × 10^5^ Huh-7 cells were infected with MOI 10 in triplicate in 6-well plates and were incubated for 12 hr, 18 hr, 24 hr, 30 hr and 36 hr prior to cell harvest using trypsin-EDTA. The cells were counted, cell numbers were normalized and lysates were prepared using RIPA buffer.

For growth curves, 4-5 × 10^5^ Huh-7 cells were infected (MOI=0.1) in triplicate in 6-well plates. Supernatants from the same wells were collected at 12 hr, 18 hr, 24 hr, 36 hr, 48 hr and 72 hr after infection. Titer was measured by plaque assay in Huh-7 .5 cells.

For flow cytometry-based curves of virus growth kinetic, 1-2 × 10^5^ Huh-7 cells were infected (MOI 0.1) in triplicates in 12-well plates. Cells were collected by trypsin-EDTA at 24 hr, 36 hr and 48h after infection. Cells were fix and permeabilized using eBioscience FoxP3 / Transcription Factor Staining Buffer Set (ThermoFisher Scientific, 00-5523-00) and stained with 0.25μg/ml of mouse monoclonal yellow fever antibody (3576) (Santa Cruz, #sc-58083) for 1 hour. After 2 washes cells were incubated for 1 hour with 0.5 μg/ml of Alexa Fluor 555 goat anti-mouse IgG (H+L) (ThermoFisher Scientific, #A21424). After 2 washes cells were resuspended in PBS and analyzed by flow cytometry using a FACS BD LSR-Fortessa.

### YFV-C antibody

Rabbit polyclonal anti-YFV-C was generated by Covance research Products. Briefly, YFV-C peptide 11-LGVNMVRRGVRSLSNK-26 was fused to KLH and 2 Rabbits were immunized with YFV-C-KLH fusion emulsified in Freund’s Complete Adjuvant. After 5 immunization boosts serum was obtained and Rabbit polyclonal anti-YFV-C was affinity purified against YFV-C immunizing peptide.

### Immunoprecipitation (IP) and western blot

HeLa cells were treated for 16 hours with Trichostatin A (4nM) and nicotinamide (100mM) prior to cell lysis. Cells were harvested by trypsin-EDTA incubation, washed twice in PBS and lysed in a modified Ripa buffer (50mM Tris pH8; 500mM NaCl; 1% NP-40; 0.5% Sodium deoxycholate; 0.1% SDS; protease inhibitor cocktail; phosphatase inhibitor cocktail; trichostatin A; nicotinamide; DNAse) and incubated for 1 hour at 4°C with rotation. Cell lysate was cleared by 16000 RPM centrifugation and the supernatant was quantified for total protein content by Bradford using BSA as standard. For IP, 1mg of lysate was incubated with 5μg of antibody for at least 16 hours. The lysate/antibody mix was then incubated with 30μL of Dynabeads Protein G (#10003D, ThermoFisher Scientific) for 1 hour at 4°C. The lysate/antibody/beads mix was then washed 5 times on the magnetic stand with modified RIPA buffer and eluted in 30μL elution buffer (50mM Glycine pH2.6; 500mM NaCl; 1X SDS loading buffer). Antibodies: BRD4 (E2A7X, Cell Signaling, #13440S), BRD3 (Bethyl, #A302-367A) and BRD2 (Bethyl, #A302-583A).

Proteins were separated in a NuPAGE 4-12% Bis-Tris Gel, transferred to PVDF membrane and probed with primary antibody for 16 hours. Antibodies utilized include anti-HA-Peroxidase High Affinity (3F10, Roche, #12013819001), anti-FLAG M2 (Sigma-Aldrich, #F1804), Lamin B (C-20, Santa Cruz, #sc-6216), and alpha-Tubulin (DM1A, Sigma-Aldrich, #T9026).

### Cell fractionation

Cells were harvested, washed twice in PBS and counted prior to lysis. Cells were incubated for 10 minutes in ice in sucrose lysis buffer (10mM Tris pH 8; 320mM sucrose; 3mM CaCl_2_; 2mM magnesium acetate; 0.1mM EDTA, 0.5% NP-40) supplemented with protease inhibitor cocktail (Sigma, #P8340). Nuclei were separated from cytoplasm by centrifugation at 500 × g at 4°C. Nuclei were washed in sucrose lysis buffer without NP-40. Nuclei were lysed in nuclei sonication buffer (50mM Tris pH8; 500mM NaCl; 1% NP-40; 10% glycerol) for 10 minutes and sonicated for 10 cycles of 30 seconds with 30 seconds intervals.

### Nuclear fractionation into chromatin and nucleoplasm

Cells were washed with PBS and lysed in a NP-40 buffer (10 mM Hepes KOH pH 7.6, 15 mM KCl, 2mM EDTA, 0.15 mM spermine, 0.5 mM spermidine, 10% glycerol, 0.3M sucrose and 0.5% NP-40) supplemented with protease and phosphatase inhibitors plus 0.5mM DTT and incubated for 10 min on ice. Nuclei were isolated by centrifugation for 10 min at 3000xg through a 30% sucrose cushion (10 mM Hepes KOH pH 7.6, 15 mM KCl, 2 mM EDTA, 0.15 mM spermine, 0.5 mM spermidine, 10% glycerol, 30% sucrose) supplemented with protease and phosphatase inhibitors plus 0.5mM DTT. Isolated nuclei were washed and resuspended in a nuclear storage buffer (10 mM Hepes KOH pH 7.6, 100 mM KCl, 0.1 mM EDTA, 0.15 mM spermine, 0.5 mM spermidine, 10% glycerol) supplemented with protease and phosphatase inhibitors plus 0.5mM DTT and incubated 30 min on ice with 1 volume 2xNUN buffer (50 mM Hepes KOH pH 7.6, 600 mM NaCl, 2M urea, 2% NP-40) supplemented with protease and phosphatase inhibitors plus 2mM DTT. The chromatin pellet was separated from the nucleoplasmic fraction by centrifugation for 30 min at 16000xg and washed with 1X NUN buffer. Chromatin-bound proteins were finally recovered by incubating the pellet for 30 min at 40°C under agitation in urea buffer (10 mM Tris pH 6.8, 5% SDS, 8M Urea, 0.5M DTT) and briefly sonicated.

### YFV-C cloning, protein expression, purification and quantification

YFV-C (nucleotides 119 to 421 relative to YFVgp1 polyprotein precursor, gene id: 1502173) was cloned from the plasmid pACNR-FLYF17D-Da plasmid into pcDNA5/FRT/TO (ThermoFisher Scientific, #V652020) and pET303/CT-His vector (ThermoFisherScientific, #K6302-03). Primers used for cloning into pcDNA5/FRT/TO were: Fw HindIII YFV-C: ggtggtAAGCTTccaccATGTCTGGTCGTAAAGCTCAGGGAAAAACCCTGGGCGTCAAT; Fw HindIII YFV-C K4RK8R: ggtggtAAGCTTccaccATGTCTGGTCGTAgAGCTCAGGGAAgAACCCTGGGCGT CAAT; Rv NotI YFV-C 3x Flag: accaccgcggccgcTCAGTCATCGTCATCCTTGTAATCGATATCATGATCTTTATAATCACCGTCATGGTCTTTGTAGTCCCCACTCCCCCCacggcgtttccttgaggacaa; Rv YFV-C HA: aacaacgcggccgcTCAAGCGTAATCTGGAACATCGTATGGGTAggatccacggcgtttccttgaggaca.

Primers used for cloning into pET303/CT-His were: Fw YFV-C XbaI: ggtggtTCTAGAatgtctggtcgtaaagctca and Rv YFV-C XhoI: accaccCTCGAGacggcgtttccttgaggaca YFV-C pET303/CT-His was expressed into BL21 Star (DE3)pLyss One Shot Chemically Competent E. Coli (ThermoFisher Scientific, #C6020-03) and bacteria grown to OD_600_ = 0.5-0.7 before addition of IPTG 0.2 mM for 2 hours. Bacteria were harvested by centrifugation at 10,000 × G for 10 minutes at room temperature followed by resuspension in histone wash buffer (50mM Tris Ph7.5; 100mM NaCl, 1mM EDTA, 1mM benzamidine). Cell were flash-frozen in liquid nitrogen and thawed in a warm water bath twice before dounce homogenization with 10-15 strokes and centrifugation for 20 minutes at 4°C at 23,000 × g. The pellet was then washed and completely homogenized in 100 mL wash buffer plus 1% Triton-X100 and centrifuged for 10 min at 4°C and 25,000 × g. The wash step was repeated at least twice, followed by two washes in wash buffer without Triton-X. The pellet was resuspended in 30 ml Protein Unfolding Buffer (20mM Tris pH7.5; 7M guanidinium; 10mM DTT) and stirred gently for 1 hour at room temperature. Insoluble material was removed by centrifugation (25,000 × g for 30 min at 20°C) prior to incubation in 1 ml of Ni-NTA agarose (ThermoFisher Scientific, #R901-01) for 2 hours, rotating at 4°C. The beads were washed 5 times in BC100 buffer (20% glycerol, 20 mM Tris-HCl, pH 7.9, at 4°C, 0.2 mM EDTA, 0.5 mM PMSF, 1.0 mM DTT, 100mM KCl) with 20mM imidazole and the bound proteins elute with BC1000 containing 400mM imidazole.

To determine YFV-C amounts per cytosol or nucleus, western blots of lysates from a known number of cells fractionated into nuclear and cytosolic fractions were performed. Varying amounts of recombinant purified YFV-C, such as; 12.g ng/lane, 25 ng/lane, 50 ng/lane, 100 ng/lane and 200 ng/lane were used to generate a standard curve. The molecular weight of YFV-C protein (residues 1-101) is 11,490 g/mole, thus 1 ng of YFV-C equals 0.0870311 picomoles (or 8.70311 × 10^-14^ moles). Using Avogadro’s number, we calculated that 1 ng of YFV-C equals 5.241 × 10^10^ molecules.

### Immunofluorescence

Following infection, cells were fixed with 4% paraformaldehyde (Electron Microscopy Sciences) for 15 min, washed with PBS and permeabilized with PBS-Triton X-100 0.1% (Sigma) for another 15 min. After a blocking step with PBS containing 5% chicken serum (Gibco), cells were incubated with anti-YFV-C and anti-YFV (Santa Cruz) primary antibodies for 45 min at room temperature. Coverslips with fixed cells were then washed with PBS and incubated with the secondary antibodies Alexa-fluor^™^ 488 chicken anti-rabbit IgG and Alexa-fluor^™^ 555 goat anti-mouse IgG (Invitrogen) for 45 min at room temperature. The coverslips were mounted onto glass slides using Prolong Gold with DAPI (Life Technologies). Images were obtained with MetaVue acquisition software (Molecular Devices) using a wide-field fluorescence microscope (Zeiss).

### Sample preparation for mass spectrometry

Proteins in the eluted sample were run in a 12% Bis-Tris gel (Life Technologies) stained with Coomassie dye. Gel bands corresponding to YFV-C protein were excised and diced into 1mm^3^ pieces taking precautions against keratin contamination. The gel pieces were washed in water followed by 100mM NaHCO_3_ and dehydrated in acetonitrile, several times before reducing the protein using 5mM dithiothreitol (Sigma) at 55°C for 45 min and alkylating in 14mM iodoacetamide (Sigma) at room temperature for 30 min in the dark. This was followed by in-gel digestion using 1:100 ratio enzyme:sample (w:w) using as enzyme sequencing grade modified trypsin (Promega, Madison, WI) overnight at 37°C. Peptides were then extracted in the solution outside the gel pieces by sequentially shrinking and swelling the gel pieces with water and acetonitrile, respectively and all the liquid was pooled for concentration in a speedvac. They peptides were subsequently de-salted using C_18_ reversed phase cartridges (Sep-Pak, Waters). After drying, peptides were resuspended in 0.1 % acetic acid for LC-MS/MS analysis and loaded onto an Easy-nLC system (Thermo Fisher Scientific, San Jose, CA, USA), coupled online with an Q Exactive mass spectrometer (Thermo Scientific). The samples were loaded on to a pulled-tip fused silica column with a 75 μm ID packed in-house with 12 cm of 3μm C_18_ resin (Reprosil-Pur C_18_-AQ). Loading buffer and buffer A was 0.1% formic acid in water, buffer B was 0.1% formic acid in acetonitrile. A gradient of 95 minutes was set for peptide elution from 2-35% buffer B. The mass spectrometer operated in positive ion mode using data dependent acquisition (DDA) with dynamic exclusion enabled (repeat count: 1, exclusion duration: 30 sec). Full MS scan (m/z 350 to 1650) was collected at a resolution of 60,000 at an AGC target value of 1×10^6^. MS2 scans of the most intense peptide ions with charge state 2-4 were performed using HCD 27 (isolation width = 2 m/z, resolution = 15,000) at an AGC target value of 5×10^4^.

### Proteomics data analysis

MS data were analyzed using Proteome Discoverer 1.4 (Thermo Scientific) for protein identification/quantification. A custom Yellow Fever Virus uniProt database was used for Mascot searches for identification of proteins and modifications. The search was performed using a precursor ion mass tolerance of 10 ppm, fragment ion mass tolerance of 0.02 Da for HCD scans, trypsin as digestion enzyme and two missed cleavages allowed. Static modifications set were carbamidomethylation of cysteine residues, and oxidation of methionine residues. Common contaminants such as keratin and trypsin were eliminated and only peptides with an FDR<1% were used to determine protein content. The mass spectrometry proteomics data have been deposited to the ProteomeXchange Consortium via the PRIDE partner repository with the dataset identifier PXD011557 and 10.6019/PXD011557.

### Protein expression and purification for structure studies

Our studies are focused on the BD1 bromodomain construct (residues 44-168) of human Brd4, termed as Brd4(BD1). The construct was expressed as a fusion protein with an N-terminal hexa-histidine tag plus yeast Sumo tag in *E. coli* strain BL21 (DE3) RIL (Stratagene, Santa Clara, CA, USA). The cells were cultured at 37°C until OD_600_ reached 0.8, following which the media was cooled to 20°C and IPTG was added to a final concentration of 0.4 mM to induce protein expression overnight at 16 °C. The cells were harvested by centrifugation at 4°C and disrupted by sonication in buffer W (500 mM NaCl, 20 mM imidazole, and 50 mM Tris-HCl, pH 7.5) supplemented with 1 mM phenylmethylsulfonyl fluoride (PMSF) protease inhibitor and 2 mM β-mercaptoethanol. After centrifugation, the supernatant was loaded onto a HisTrap FF column (GE Healthcare, Little Chalfont, Buckinghamshire, UK). After extensive washing by buffer W, the target protein was eluted with buffer W supplemented with 300 mM imidazole. The hexa-histidine-Sumo tag was cleaved by Ulp1 protease and removed by further elution through a HisTrap FF column. Individual pooled target proteins were further purified by passage through a Hiload 16/60 Superdex 200 column (GE Healthcare) with buffer E (150 mM NaCl, 10 mM Tris-HCl, pH 7.5, and 1 mM DTT).

### Crystallization and data collection

Brd4(BD1) was collected by pooling Hiload 16/60 Superdex 200 (GE Healthcare) gel filtration chromatographic peak fractions to a concentration of 25 mg/mL for crystallization trials. Before crystal screening, the Brd4(BD1) protein was mixed with YFV(1-10) K4ac, K8ac and K4ac/K8ac peptides in a molar ratio of 1:3 at 4°C for 2 h. Crystallization of the Brd4(BD1)-YFV(1-10) acetylated peptides were carried out at 20 °C using the hanging drop vapor diffusion method by mixing 0.2 μL protein sample at a concentration of 12 mg/ml and 0.2 μL reservoir solution and equilibrating against 80 μL reservoir solution. Crystals of Brd4(BD1) with YFV-C (1-10) K4ac peptide were grown from a solution containing 0.2 M sodium chloride (QIAGEN), 0.1 M Tris (pH 7.0) (QIAGEN), 30% PEG 3000 (QIAGEN) at 20 °C using the hanging drop vapor diffusion method. Crystals of Brd4(BD1) with YFV-C (1-10) K8ac peptide were grown from a solution containing 0.1 M HEPES (pH 7.0) (QIAGEN), 30% PEG 6000 (QIAGEN) at 20 °C using the hanging drop vapor diffusion method. Crystals of Brd4(BD1) with YFV-C (1-10) K4ac/K8ac peptide were grown from a solution containing 0.1 M HEPES (pH 7.0) (QIAGEN), 30% PEG 6000 (QIAGEN) at 20 °C using the hanging drop vapor diffusion method. For data collection, crystals were flash frozen (100 °K) in the above reservoir solution supplemented with 10% glycerol (QIAGEN). The diffraction data set for complexes of Brd4(BD1) with YFV-C (1-10) K4ac, YFV-C (1-10)K8ac and YFV-C (1-10) K4ac/K8ac peptides were collected to 1.5 Å, 1.4 Å and 1.4 Å resolution, respectively. All data sets were collected at wavelength 0.979 Å on NE-CAT beam line 24ID-C at the Advanced Photon Source (APS), Argonne National Laboratory. All the data were processed with the program HKL2000^5^. The statistics of the diffraction data are summarized in Supplementary Table S1.

### Structure determination and refinement

All three structures of complexes of Brd4(BD1) with bound YFV-C (1-10) acetylated peptides were solved using molecular replacement method as implemented in the program PHENIX^6^ based on the Brd4(BD1) fold in the Brd4(BD1)-H4 K5ac/K8ac peptide structure (PDB: 3UVW). All the molecular graphics were generated with the program Pymol (DeLano Scientific).

### ChIP-Sequencing

Huh-7 cells, Huh-7 YFV-C and Huh-7 YFV-C K4RK8R cells were grown at 37°C and 5% CO_2_ for 48 hours in complete culture medium with 10ug/ml doxycycline. Cells were harvested by trypsinization, pelleted and resuspended in PBS at a final concentration of 1 × 10^6^ cells/ml. Cells were fixed with 1% formaldehyde (methanol-free) for 10 minutes. The formaldehyde was quenched for 5 minutes by addition of glycine to a final concentration of 0.125M. Cells were washed twice with ice cold PBS and then used for ChIP as previously described^7^. In brief, Chromatin was sonicated to an average size of 200-600bp. BRD4-associated chromatin fragments were IP-ed ON at 4C using a rabbit monoclonal anti-BRD4 antibody (Cell Signaling, #134403S) coupled to anti-rabbit M-280 Dynabeads (ThermoFisher Scientific, #11204D). After 12 hours incubation at 4°C the beads were washed with RIPA wash buffer with 100mM LiCl and TE wash buffer as previously described^7^. After elution, decrosslinking and treatment with RNase and proteinase K the IP-ed DNA was purified and used for library generation.

ChIP-Seq libraries were generated as previously described^7^. In brief, the DNA was blunt-ended, an “A”-overhang was added and Illumina Paired-End adaptors were ligated to the DNA fragments. ChIP-Seq library amplification was performed using Illumina primers 1.0 and 2.0. Libraries were sequenced on the HighSeq2500 platform for 50 cycles. The raw data for all ChIP-Seq libraries were processed following the ENCODE pipeline. Adaptor sequences were removed using Trimmomatic version 0.36 using default settings (ILLUMINACLIP:adapter.fa:2:30:7 SLIDINGWINDOW:3:15 MINLEN:36). The trimmed sequencing data were aligned to the human genome (GRCh37/hg19) using Bowtie2-2.2.5 using default settings. Unmapped reads, QC-failed reads and reads with low mapping quality (-F 0×4 –q 10) were removed to generate the final bam files.

Peak calling was performed using macs2 version 2.1.1. The extsize of each library was predicted using “macs2 predictd”. Peak calling was performed for each replicate individually with respect to the matching input control using “macs2 callpeak” with the parameters (-g hs --broad --nomodel--extsize 75). Transcriptional start site (TSS)peaks were defined as peaks located within 1kb from the TSS. Principle component analysis (PCA) was performed using the R function prcomp on the log normalized read counts for the BRD4 binding atlas.

Differential binding analysis was performed using the R-package DESeq2 for Huh-7 - Tet on YFV-C versus Huh-7. Peaks with a Benjamini & Hochberg (BH) adjust P-value < 0.05 were considered to be significantly differentially associated with BRD4. BRD4 ChIP-seq clustering was generated using k-means with the pheatmap R-package (v1.0.12).

### RNA-Sequencing

RNA was isolated using the RNeasy mini kit (Qiagen, #74106) and quality was analyzed on the Agilent 2100 Bioanalyzer and all samples had RNA integrity number higher than 8.0. 100 ng of total RNA was used to generate RNA-Seq libraries using Illumina TruSeq stranded mRNA LT kit (Cat# RS-122-2101). Libraries prepared with unique barcodes were pooled at equal molar ratios. The pool was denatured and sequenced on Illumina NextSeq 500 sequencer using high output V2 reagents and NextSeq Control Software v1.4 to generate 75 bp single reads, following manufactures protocol (Cat# 15048776 Rev.E).

Sequencing data were aligned to the human genome (GRCh37/hg19) using the STAR alignment tool. The aligned reads were filtered for uniquely aligned reads with MAQ > 10. The aligned RNA-Seq reads were counted using hg19 RefSeq annotation and the SummarizeOverlaps function from the R-package GenomicAlignments with IntersectionNotEmpty mode. Around 20 million reads were counted for each sequencing library. Principle component analysis (PCA) was performed using the R function prcomp on log normalized read counts. Differential expression analysis was performed using the R-package DESeq2 for two comparisons Huh-7-YFV-C versus Huh-7. Genes with BH adjust P-value < 0.001 were considered to be differentially expressed.

GSEA analysis was performed using the R-package clusterProfiler (v3.12.0)^9^, using 10^6^ permutations of gene ontology biological processes with a minimum and maximun geneset size of 50 and 250, respectively. RNAseq clustering was generated using k-means with the pheatmap R-package (v1.0.12).

### ATAC-Sequencing

The ATAC–seq assay was performed on 75,000 Huh-7, Huh-7 YFVC and Huh-7 YFVC K4RK8R cells as previously described^10,11^. In brief, cells were lysed in ATAC lysis buffer for 5 min and then transposed with TN5 transposase (Illumina) for 30 min. Samples were barcoded and the sequencing library was prepared according to manufacturer’s guidelines (Illumina) and sequenced on an Illumina HiSeq 2500. ATAC-seq reads were aligned to the hg19 genome from the Bsgenome.Hsapiens.UCSC.hg19 Bioconductor package (version 1.4.0) using Rsubread’s align method in paired-end mode with fragments between 1 to 5000 base-pairs considered properly paired^12^. Normalized, fragment signal bigWigs were created using the rtracklayer package. Peak calls were made with MACS2 software in BAMPE mode^13^. Differential ATAC-seq signal is identified using the DESeq2 package^14^.

### Affinity measurements

Surface Plasmon Resonance (SPR) binding experiments were carried out on a ProteOn XPR36 6×6 channels protein interaction array system (Bio-Rad). Proteins BRD4-BD1 an BRD4-BD2 were diluted to a final concentration of 50μg/ml in HEPES buffer 25 mM pH 7.0 and immobilized to a GLM chip using amine coupling. The immobilization level was about 2,000 response units (RU). Analytes (YFV-C and histone H4 peptides) were diluted in 10 mM HEPES pH 7.0, 150 mM NaCl, 5 mM MgCl_2_ and 0.01% Tween-20 and serial dilutions were injected over the BRD4-BD1 an BRD4-BD2 coated chip at a flow rate of 100 μl/min for 60 s and were subsequently allowed to dissociate for 600 s. The chip surface was regenerated by sequential injections of buffer 10 mM HEPES pH 7.0, 150 mM NaCl, 5 mM MgCl_2_ and 0.01% Tween-20. Reproducibility was assessed with two additional experiments performed on independent chips. The experiments were carried out at 25°C. Data was normalized by interspot and buffer subtraction. Data was analyzed using equilibrium between 45 and 50 sec.

## Acknowledgements

The work was supported in part by grant AI124690 (to C.M.R.), NIH grants AI118891, GM110174 and CA196539 (to B.A.G.), NIH grant CA234561 (to R.G.R), Human Frontier Science Program Long-Term Fellowship (LT001083/2014) (to A.G.) and CAPES/Brazil (to L.A.C.).

## Author contributions

Diego Mourao, Shoudeng Chen and Uwe Schaefer contributed equally to this work.

Margaret R. MacDonald, Dinshaw Patel and Alexander Tarakhovsky contributed equally to this work.

D.M. and A.T. conceived the project. D.M performed most of the experiments, analyzed, interpreted data and prepared the manuscript. A.G. performed experiments to determined YFV-C interaction with chromatin and A.G. and R.G.R. conceived and analyzed YFV-C interaction with chromatin. S.C. and D.P. performed, analyzed and interpreted crystal structure data. U.S. performed ChIP-seq experiments and virus infection experiments. L.B. generated mutant YFV, performed virus infection and virus quantification assays. L.A.C. performed and analyzed ATAC-seq experiments and immunofluorescence microscopy. C.A. performed and analyzed SPR data. B.D and H.M. performed and analyzed MS/MS data YFV-C acetylation from ectopic expressing cells. T.C. and M.P. analyzed and advised on RNA-seq, ATAC-seq and ChIP-seq data. N.B and B.G. performed and analyzed MS/MS data on YFV-C acetylation from infected cells. G.J, I.R, R.P. conceived, analyzed and interpreted data with BET inhibitor and pilot MS/MS experiments. L.B., M.R.M and C.M.R. conceived and analyzed YFV generation, infection and phenotyping. D.M. wrote the manuscript with input from A.T., D.P., M.R.M, C.R.M. and other authors.

## References

1. Davey, N. E., Travé, G. & Gibson, T. J. How viruses hijack cell regulation. Trends Biochem Sci 36, 159 169 (2011).

2. Daugherty, M. D. & Malik, H. S. Rules of engagement: molecular insights from host-virus arms races. Annu Rev Genet 46, 677 700 (2012).

3. Haerty, W. & Ponting, C. P. No gene in the genome makes sense except in the light of evolution. Annu Rev Genom Hum G 15, 71 92 (2014).

4. Malik, H. S. & Henikoff, S. Phylogenomics of the nucleosome. Nat Struct Mol Biology 10, 882 891 (2003).

5. Lindenbach, B. D. & Rice, C. M. Molecular biology of flaviviruses. Advances in virus research 59, 23 61 (2003).

6. Mukhopadhyay, S., Kuhn, R. J. & Rossmann, M. G. A structural perspective of the flavivirus life cycle. Nat Rev Microbiol 3, 13 22 (2005).

7. Byk, L. A. & Gamarnik, A. V. Properties and Functions of the Dengue Virus Capsid Protein. Ann Rev Virol 3, 263 281 (2016).

8. Colpitts, T., Barthel, S., Wang, P. & Fikrig, E. Dengue Virus Capsid Protein Binds Histones and Inhibits Nucleosome Formation in Human Liver Cells. Plos One 6, e24365 (2011).

9. Wuarin, J. & Schibler, U. Physical isolation of nascent RNA chains transcribed by RNA polymerase II: evidence for cotranscriptional splicing. Mol Cell Biol 14, 7219 7225 (1994).

10. Ramage, H. R. et al. A Combined Proteomics/Genomics Approach Links Hepatitis C Virus Infection with Nonsense-Mediated mRNA Decay. Mol Cell 57, 329–340 (2015).

11. Li, M. et al. Identification of antiviral roles for the exon–junction complex and nonsense-mediated decay in flaviviral infection. Nat Microbiol 4, 985–995 (2019).

12. Shah, P. S. et al. Comparative Flavivirus-Host Protein Interaction Mapping Reveals Mechanisms of Dengue and Zika Virus Pathogenesis. Cell 175, 1931 1945.e18 (2018).

13. Carlin, A. F. et al. Deconvolution of pro- and antiviral genomic responses in Zika virus-infected and bystander macrophages. Proc National Acad Sci 115, 201807690 (2018).

14. Pickett, B. E. et al. ViPR: an open bioinformatics database and analysis resource for virology research. Nucleic Acids Res 40, D593–D598 (2012).

15. Jones, C. T. et al. Flavivirus capsid is a dimeric alpha-helical protein. J Virol 77, 7143 7149 (2003).

16. Shahbazian, M. D. & Grunstein, M. Functions of site-specific histone acetylation and deacetylation. Annu Rev Biochem 76, 75 100 (2007).

17. Belkina, A. C. & Denis, G. V. BET domain co-regulators in obesity, inflammation and cancer. Nat Rev Cancer 12, 1 13 (2012).

18. Filippakopoulos, P. & Knapp, S. The bromodomain interaction module. Febs Lett 586, 2692 2704 (2012).

19. Dhalluin, C. et al. Structure and ligand of a histone acetyltransferase bromodomain. Nature 399, 491 496 (1999).

20. Filippakopoulos, P. et al. Histone Recognition and Large-Scale Structural Analysis of the Human Bromodomain Family. Cell 149, 214 231 (2012).

21. Wisniewski, J. R., Vildhede, A., Norén, A. & Artursson, P. In-depth quantitative analysis and comparison of the human hepatocyte and hepatoma cell line HepG2 proteomes. J Proteomics 136, 234 247 (2016).

22. Lachmann, A. et al. ChEA: transcription factor regulation inferred from integrating genomewide ChIP-X experiments. Bioinformatics 26, 2438 2444 (2010).

23. Patkar, C. G., Jones, C. T., Chang, Y., Warrier, R. & Kuhn, R. J. Functional requirements of the yellow fever virus capsid protein. J Virol 81, 6471 6481 (2007).

24. Rice, C. M. et al. Nucleotide sequence of yellow fever virus: implications for flavivirus gene expression and evolution. Science 229, 726 733 (1985).

25. Hahn, C. S. et al. Conserved elements in the 3′ untranslated region of flavivirus RNAs and potential cyclization sequences. J Mol Biol 198, 33–41 (1987).

26. Nicodeme, E. et al. Suppression of inflammation by a synthetic histone mimic. Nature 468, 1119 1123 (2010).

## Methods References

1. Bredenbeek, P. J. et al. A stable full-length yellow fever virus cDNA clone and the role of conserved RNA elements in flavivirus replication. J Gen Virol 84, 1261–1268 (2003).

2. Amberg, S. M. & Rice, C. M. Mutagenesis of the NS2B-NS3-mediated cleavage site in the flavivirus capsid protein demonstrates a requirement for coordinated processing. Journal of virology 73, 8083 8094 (1999).

3. Yi, Z. et al. Identification and characterization of the host protein DNAJC14 as a broadly active flavivirus replication modulator. Plos Pathog 7, e1001255 (2011).

4. Kümmerer, B. M. & Rice, C. M. Mutations in the yellow fever virus nonstructural protein NS2A selectively block production of infectious particles. Journal of virology 76, 4773 4784 (2002).

5. Otwinowski, Z. & Minor, W. Processing of X-ray diffraction data collected in oscillation mode. Methods Enzymol 276, 307–26 (1997).

6. Adams, P. D. et al. PHENIX: a comprehensive Python-based system for macromolecular structure solution. 539–547 (2012) doi:10.1107/97809553602060000865.

7. Nicodeme, E. et al. Suppression of inflammation by a synthetic histone mimic. Nature 468, 1119 1123 (2010).

8. Santiago, I. de & Carroll, T. Methods in Molecular Biology. Methods Mol Biology Clifton N J 1689, 195–226 (2017).

9. Yu, G., Wang, L.-G., Han, Y. & He, Q.-Y. clusterProfiler: an R package for comparing biological themes among gene clusters. Omics J Integr Biology 16, 284–7 (2012).

10. Buenrostro, J. D., Wu, B., Chang, H. Y. & Greenleaf, W. J. ATAC-seq: A Method for Assaying Chromatin Accessibility Genome-Wide. Curr Protoc Mol Biology Ed Frederick M Ausubel Et Al 109, 21.29.1–9 (2015).

11. Buenrostro, J. D., Giresi, P. G., Zaba, L. C., Chang, H. Y. & Greenleaf, W. J. Transposition of native chromatin for fast and sensitive epigenomic profiling of open chromatin, DNA-binding proteins and nucleosome position. Nat Methods 10, 1213 1218 (2013).

12. Liao, Y., Smyth, G. K. & Shi, W. The Subread aligner: fast, accurate and scalable read mapping by seed-and-vote. Nucleic Acids Res 41, e108 (2013).

13. Feng, J., Liu, T., Qin, B., Zhang, Y. & Liu, X. S. Identifying ChIP-seq enrichment using MACS. Nat Protoc 7, 1728–40 (2012).

14. Ross-Innes, C. S. et al. Differential oestrogen receptor binding is associated with clinical outcome in breast cancer. Nature 481, 389–93 (2012).

